# Domain-General Brain Networks Support Language Development

**DOI:** 10.1101/2025.07.10.663607

**Authors:** Zihang Zhou, Xiaohong Yang, Ping Ju, Mengjie Rong, Elizabeth Jefferies, Xi-Nian Zuo

## Abstract

Understanding the neural basis of verbal intelligence across development requires disentangling the contributions of domain-general and language-selective brain systems. Although language is often considered a domain-specific function, complex language tasks also engage domain-general networks, such as the Default-Mode (DM) and Multiple-Demand (MD) systems. Yet how these systems contribute to the maturation of verbal competence remains poorly understood. Here, we examined this question using gray matter volume measures in an accelerated longitudinal dataset of children and adolescents from Beijing (N = 170), using the Verbal Comprehension Index (VCI) from the Wechsler Intelligence Scale as a benchmark for verbal abilities. We observed that individual differences in VCI were more strongly associated with structural maturation of domain-general networks (DM and MD) than with the language-selective network, and that these effects varied with age. Targeted validation in an independent cohort from Chongqing (N = 150) confirmed significant contributions of domain-general networks in adolescence (13–15 years), highlighting the robustness of these developmental effects. These findings suggest that domain-general cortical systems play a critical and previously underappreciated role in the emergence of verbal intelligence during adolescence, with implications for understanding how large-scale brain networks support the development of abstract verbal reasoning.

## Main

Language competence is a hallmark of human cognition, yet the neural systems that support it—particularly during development—remain a matter of ongoing debate. While much is known about the mature architecture of language processing (e.g., Braga et al, 2020; Caplan, 1994; Fedorenko and Thompson Schill, 2014; Friederici, 2011; Friederici and Gierhan, 2013; Friederici and Wartenburger, 2010; Hickok, 2009; Price, 1998; Turker et al, 2023), less is understood about how domain-specific and domain-general networks shape language abilities across childhood and adolescence. A key theoretical tension persists between two frameworks: (1) the domain-specific view (Brauer and Friederici, 2007; Friederici, 2011), which proposes that language is a dedicated cognitive domain supported by a specialized neural network; and (2) the domain-general view (Fedorenko et al, 2024a); for reviews, see (Fedorenko and Varley, 2016), which argues that although core language functions are distinct from higher-order cognition, the application of language in complex contexts recruits domain-general systems such as the Default-Mode (DM) and Multiple-Demand (MD) networks. Recent findings further complicate this debate by demonstrating that while language ability dissociates from general intelligence (Fedorenko and Varley, 2016; Mahowald et al, 2024), complex language tasks consistently engage domain-general networks (Hu et al, 2023; Sherafati et al, 2022). This paradox highlights the need to clarify how domain-specific and domain-general networks jointly support language competence. The current study addresses this gap by investigating how morphological changes in both network types contribute to language competence through development, testing competing predictions of the two frameworks.

Language processing recruits a domain-specific language-selective network, which includes a set of left-lateralized frontal and temporal regions (Binder et al, 1997; Fedorenko et al, 2010, 2011; Fedorenko and Thompson Schill, 2014; Friederici, 2011; Scott et al, 2017; Vigneau et al, 2006). Functional magnetic resonance imaging (fMRI) studies consistently demonstrate a high degree of functional specificity within these brain regions, emphasizing their role in language processing with minimal neural overlap with other cognitive processes (Blank and Fedorenko, 2017; Fedorenko and Shain, 2021; Fedorenko et al, 2013; MacGregor et al, 2022; Mineroff et al, 2018; Shain et al, 2020, 2022, 2023). Further investigations have validated the functional selectivity of this language-selective network across different languages (Malik-Moraleda et al, 2022), highlighting its robustness. Yet recent studies also recognize that language processing extends beyond statistical regularities necessary for language comprehension and production – instead, language functions as a tool that enables individuals to achieve goals and build relationships in real-world situations. This functional linguistic capacity may depend on additional cognitive capabilities (for reviews, see Mahowald et al, 2024).

Activation has been identified in brain networks that fall outside of language-selective regions, particularly in the MD network (e.g., Duncan, 2010; Duncan and Owen, 2000; Hsu et al, 2017); for reviews, see (Fedorenko et al, 2024a; Ryskin and Nieuwland, 2023) and the DM network (Buckner et al, 2008; Mattheiss et al, 2018). The MD network shows a well-documented response in demanding cognitive tasks and is thought to support response inhibition, working memory and goal-directed behavior (Niendam et al, 2012; Stiers et al, 2010). Engagement of the MD network has been reported in relation to diverse aspects of language tasks, including morphosyntax (Hsu et al, 2017; Newhart et al, 2012; Peelle et al, 2010), semantics (Davey et al, 2016; Hasson et al, 2018; Novick et al, 2005; Whitney et al, 2012), speech perception (MacGregor et al, 2022; Sherafati et al, 2022), and discourse processing (Yang et al, 2023); this network is recruited when we answer questions (Diachek et al, 2020; Hu et al, 2023). Recent evidence also emphasizes the importance of the DM network, especially the dorsomedial subsystem, in discourse comprehension (Yang et al, 2023) and in representing goal-related semantic information (Liu et al, 2022; Wang et al, 2021); for reviews, see (Menon, 2023), suggesting its essential role in constructing mental representations of coherent narrative. These findings together show that language processing is not solely dependent on the language-selective network but also derives support from domain-general cognitive networks. This highlights the interconnected and distributed nature of the neural basis of language and its application in real-world contexts.

Yet the debate about whether language processing relies on domain-general cognitive resources remains unresolved. Some evidence suggests that the core contribution of these networks to language may be overstated, as language comprehension predominantly engages the language-selective network, with minimal reliance on domain-general brain networks (Blank and Fedorenko, 2017; Branzi and Lambon Ralph, 2023; Diachek et al, 2020); for reviews, see (Campbell and Tyler, 2018; Ryskin and Nieuwland, 2023). The primary debate centers on the MD network’s poor tracking of core linguistic computations, such as lexical access, syntactic parsing and semantic composition, across individuals (e.g., Blank and Fedorenko, 2017; Diachek et al, 2020; Wehbe et al, 2021); for reviews, see (Fedorenko and Shain, 2021). These findings challenge the assertion that the MD network plays a core role in receptive language functions (Clercq et al, 2024). By this view, general-purpose cognitive systems like the MD network and DM network are only indispensable when handling extraneous task demands and difficulties during comprehension. Critically, neuroimaging and aphasia research reveals a clear dissociation between language and complex thought in adults (for reviews, see Fedorenko and Varley, 2016), supporting a limited role of domain-general networks in language while addressing the centrality of linguistic representations in high-level language processing.

Conflicting views on the role of domain-general systems in language may stem from how language processing is defined and measured. While core linguistic operations, such as syntactic parsing or single-word comprehension, may proceed independently of domain-general systems during passive tasks, real-world language use typically involves integrating contextual, pragmatic, and social information, placing heavier demands on higher-order cognitive resources (Ryskin and Nieuwland, 2023). To better capture these demands, the present study employed the Wechsler Verbal Comprehension Index, a standardized measure of verbal intelligence that reflects broader language competence. Unlike tasks using artificially constrained stimuli or passive comprehension paradigms, this test assesses expressive vocabulary, verbal reasoning, and acquired knowledge, and has been shown to predict real-life communicative abilities with greater ecological validity (Kuehnel et al, 2019; Smith et al, 2005; Whipple Drozdick and Munro Cullum, 2011). By using the Wechsler Verbal Comprehension Index, we aimed to investigate how domain-general systems support the development of linguistic abilities that extend beyond narrow definitions of language processing, providing insight into the neural bases of functional language use in everyday life.

Children and adolescents, with their relatively lower language proficiency, may depend more heavily on domain-general networks to compensate for limitations in the language-selective network (Rosselli et al, 2014). Adults show better performance on tests of phonology, vocabulary, and grammar, compared with children (Berken et al, 2017; Nayak et al, 2022; Rosselli et al, 2014). Children and adolescents may recruit additional resources from domain-general networks to compensate for these language immaturities; therefore, these younger age groups may rely more heavily on domain-general cognition. However, prior research has predominantly focused on adults, limiting our understanding of how large-scale brain networks collaborate to support earlier language development. In addition, domain-general networks may support multiple cognitive domains early in development (Hiersche et al, 2024), including the use of language for critical thinking, problem solving, and social cooperation—skills that are essential for success in adulthood (Baldo et al, 2005; Bloom and Keil, 2001; Maynard and Turowetz, 2013; Turkstra et al, 2017). In this study, we ask how domain-general networks vary in their cooperation with the language network in children and adolescents to gain a more comprehensive understanding of how the brain networks supporting language evolve over time.

To characterize the involvement of language-specific and domain-general networks in the development of language competence, we analyzed two accelerated longitudinal datasets: the devCCNP-PEK sample from Beijing (n= 170) in our primary analysis and the devCCNP-CKG sample from Chongqing (n= 150) for the validation of key findings. We examined associations between cortical morphology and overall language competence across childhood and adolescence. Specifically, we measured gray matter volumes within the language, MD, and DM networks and utilized the Verbal Comprehension Index (along with the scores of its subtests) as the indicator of overall language competence. Next, we modeled developmental trajectories of both network volumes and the Verbal Comprehension Index, testing their age-related dynamic associations. Finally, we assessed the unique and relative contribution of the language, multiple demand, and default mode networks in predicting variance in overall language competence throughout childhood and adolescence.

Following previous reports on longitudinal changes in cortical structures and language abilities (Mattheiss et al, 2018; Wang et al, 2021), we expected that both brain networks structures and overall language competence will exhibit age-related development, and that these effects will be associated. Given the ongoing theoretical debate between domain-specific (Brauer and Friederici, 2007; Friederici, 2011) and domain-general views (Fedorenko et al, 2024a); (for reviews, see Fedorenko and Varley, 2016), we anticipated two competing patterns of results regarding network contributions to language development. If language development is bootstrapped by domain-general networks, language competence might be more strongly related to the maturation of those networks. Alternatively, if intelligent thought is largely independent of language, language competence might be expected to be linked more specifically to the maturation of the language network.

## Materials and Methods

### Participants

Data from 320 participants in the Chinese Color Nest Project (CCNP, 2013-2032) were used in the present study, which comprised two samples: the devCCNP-PEK Sample (N = 170), collected in the Chaoyang District of Beijing, and the devCCNP-CKG Sample (N = 150), collected in the Beibei District of Chongqing (Fan et al, 2023). The Chongqing dataset was used to test the generalizability of significant age windows identified in the Beijing dataset, particularly during adolescence, to ensure robustness of findings across different populations. The CCNP employed an accelerated longitudinal design (ALD) representing a fusion of longitudinal and cross-sectional elements. This approach allows for the measurement of within-subject changes over time while also encompassing a wide age range in a manageable time frame (Thompson et al, 2011). Originally, 480 typically developing children aged 6-18 years (20 boys and 20 girls in each cohort, 12 age cohorts in total) were invited to participate in three assessment times of data collection, referred to as three waves of visits (Liu et al, 2021). The interval between two neighboring waves was approximately 15 months to balance the season effects (Fan et al, 2023). Visual inspection was carried out to eliminate scans exhibiting head-motion artifacts or those with structural abnormalities, resulting in the exclusion of 97 scans from 80 participants. Participants aged 16 and 17 years (4 in the devCCNP-PEK Sample, 27 in the devCCNP-CKG Sample) were excluded because the small number in the two cohorts provided insufficient statistical power for reliable developmental modeling, restricting the study to ages 6–15 years for robust time-varying and trajectory analyses. Additionally, 49 participants without test scores were excluded. After exclusion, a total of 585 scans from 320 participants (150 females) were included in follow-up analyses (see Table 1). Detailed sample distributions for both cohorts are provided in Supplementary Materials A (Figure S1).

**Table 1.** Demographics and Behavioral Statistics for Two Samples across all Three Waves.

|  | Wave 1 | Wave 2 | Wave 3 | Total (Scans) |
| --- | --- | --- | --- | --- |
| **The PEK Sample** |  |  |  |  |
| N(M/F)<sup>1</sup> | 170 (94/76) | 65 (38/27) | 20 (12/8) | 255 (144/111) |
| Age | 9.08 (2.19) | 10.11 (2.09) | 11.55 (1.88) | 9.54 (2.26) |
| Range | 6 - 15 | 7 - 15 | 8 - 15 | 6 - 15 |
| Vocabulary | 13.21 (3.02) | 14.38 (2.96) | 15.20 (3.38) | 13.67 (3.10) |
| Similarities | 14.26 (2.51) | 14.92 (2.29) | 15.70 (2.13) | 14.54 (2.46) |
| Comprehension | 13.52 (3.90) | 14.8 (4.17) | 16.25 (4.12) | 14.06 (4.06) |
| VCI<sup>2</sup> | 121.73 (17.41) | 128.45 (16.83) | 135.85 (19.15) | 124.55 (17.88) |
| **The CKG Sample** |  |  |  |  |
| N(M/F)<sup>1</sup> | 150 (76/74) | 124 (66/58) | 56 (31/25) | 330 (173/157) |
| Age | 10.51 (2.47) | 11.06 (2.07) | 11.75 (1.82) | 10.93 (2.26) |
| Range | 6 - 15 | 7 - 15 | 9 - 15 | 6 - 15 |
| Vocabulary | 13.03 (2.66) | 12.74(2.34) | 11.98 (2.32) | 12.74 (2.51) |
| Similarities | 12.72 (2.80) | 13.15 (2.42) | 12.89 (2.83) | 12.91 (2.67) |
| Comprehension | 13.75 (3.82) | 14.17 (3.76) | 12.32 (3.74) | 13.67 (3.83) |
| VCI<sup>2</sup> | 119.25 (16.35) | 120.37 (15.00) | 114.55 (15.49) | 118.87 (15.79) |

<sup>1</sup>N (M/F) indicates the total number of participants in each wave (and the discrimination based on sex).
<sup>2</sup>VCI: the Verbal Comprehension Index. For the rows of age, Vocabulary, Similarities, Comprehension, and VCI in the table, the data displayed represent the mean and standard error (in parentheses).

### Overall Language Competence

The Wechsler Intelligence Scale for Children-IV-Chinese Version (WISC-IV) was administered to measure cognitive ability (Wechsler, 2003) in the Project. The present study utilized the Verbal Comprehension Index from WISC-IV as a standardized language measure that may serve as indicator of overall language competence. Specifically, participants were presented with three core subtests. The Vocabulary subtest examines participants’ knowledge of acquired word concepts through word definition. The Similarities subtest assesses participants’ ability to identify the commonality between two given objects or concepts with a brief description. This task is designed to measure participants’ abilities in the formation of language concepts, and in abstract reasoning.

The Comprehension subtest was employed to test participants’ verbal reasoning and conceptualization, and their ability to demonstrate practical knowledge and judgement according to their understanding of real-life situations and social norms (Weiss et al, 2015). Detailed information regarding the contents and formats of these tests is provided in Figure 1. The subtests were measured one-on-one, and participants’ responses were manually scored (raw scores) by experimenters and then normalized (normative scores) to produce the Verbal Comprehension Index. All experimenters completed training for WISC-IV scoring. Descriptive statistics for behavioral measures for the whole group across all waves are listed in Table 1.

**Fig. 1.**
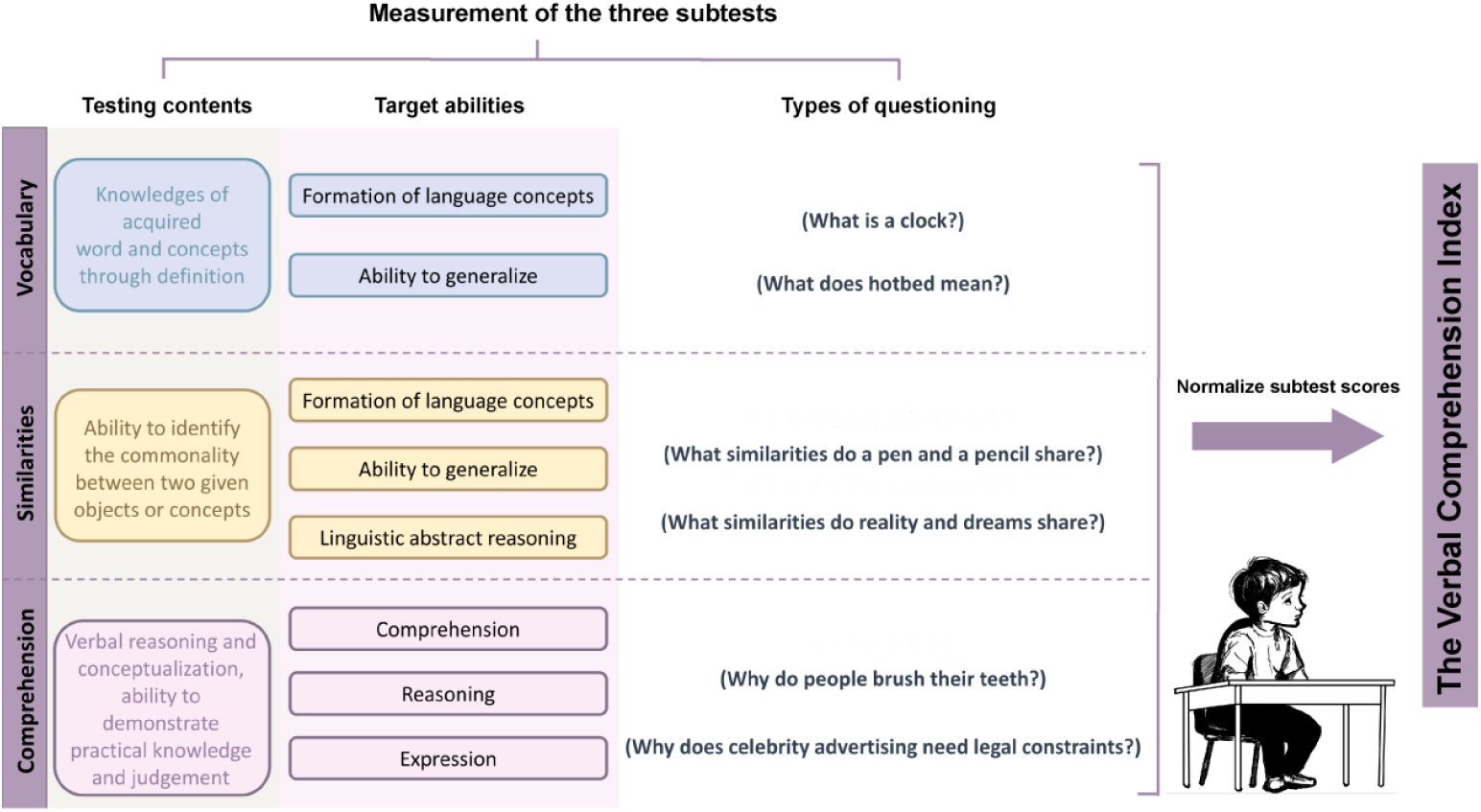
The Verbal Comprehension Index and its Subtest Measurements.

### MRI Data Acquisition

T1-weighted MRI data for the PEK Sample were collected on a 3.0-T GE Discovery MR750 scanner with an eight channel array head coil at CAS Institute of Psychology. Images were collected using a 3D T1-weighted spoiled gradient recalled (SPGR) echo sequence with the following parameters: T1 = 450 ms; TR = 6.70 ms; TE = 2.90 ms; flip angle = 12°; image matrix = 256×256; FOV = 256×256 mm^2^; 176 sagittal slices; spatial resolution = 1×1×1 mm^3^. Data in the CKG Sample were collected using a 3.0-T Siemens Trio MRI scanner with a 12 channel array head coil at the Center for Brain Imaging, Southwest University. The imaging protocol employed a 3D magnetization-prepared rapid gradient-echo (MP-RAGE) sequence with the following parameters: T1 = 900 ms; TR = 2600 ms; TE = 3.02 ms; flip angle = 8°; image matrix = 256×256; FOV = 256×256 mm^2^; 176 sagittal slices; spatial resolution = 1×1×1 mm^3^. Each scanning site maintained identical protocols across all waves, while inter-site variations were preserved with parameter optimization ensuring comparable spatial and temporal resolutions.

### MRI Data Processing

#### MRI Preprocessing

Structural MRI data were preprocessed using the Computational Anatomy Toolbox (CAT12; http://dbm.neuro.uni-jena.de/cat12) implemented in SPM 12 (Wellcome Centre of Imaging Neuroscience, Institute of Neurology, UCL, London, UK; http://www.fil.ion.ucl.ac.uk/spm). T1-weighted images were initially segmented into gray matter (GM), white matter (WM), and cerebral spinal fluid (CSF) tissues classes using the standard Tissue Probability Maps (TPMs). The segmented images were then warped to the Geodesic Shooting template (Ashburner and Friston, 2011) and normalized to the Montreal Neurological Institute (MNI) space with default settings in 1.5 mm cubic resolution. Data quality was measured by the weighted average of image quality ratings (IQR) generated by CAT12, defined by spatial resolution, Brain Web Phantom (BWP) noise, and BWP bias. Images rated below the satisfactory threshold (C; 0.75) were visually checked and manually excluded from the dataset.

#### Gray Matter Volume Extraction

The “Estimate mean values inside ROI” function implemented in CAT12 was applied to extract gray matter volume values from the three networks for each participant and for each scanning wave. The three networks of interest were the language network, the MD network and the DM network. Volumes of these large-scale networks were determined using masks obtained from a recent study by Kong et al (2021) on individual-specific 400-region parcellation from fMRI data, as shown in Figure 2. Given the sophisticated process of executive functioning, the MD network has been defined by convergent activation across multiple cognitive tasks in fMRI (Duncan, 2010). This highlights co-activation of several brain networks including the ventral attention network (Japee et al, 2015), the salience network Seeley et al (2007), the dorsal attention network (Corbetta et al, 2008), and the cognitive control network (Cole and Schneider, 2007). Therefore, we integrated seven networks including Salience/VenAttn A, Salience/VenAttn B, Control A, Control B, Control C, Dorsal Attention A and Dorsal Attention B from (Kong et al, 2021) in our definition of the large-scale MD network. We also conducted exploratory analyses using the functionally-defined MD regions provided by Fedorenko et al (2013). Our main findings were successfully replicated; detailed results are presented in Supplementary Materials C. Gray matter volume in the DM network was determined by the combination of the three DM sub-networks (A, B and C) (Buckner et al, 2008; Yeo et al, 2011). For more details about the areal-level cortical parcellation network used in the present study for network-level gray matter volume extraction, see (Kong et al, 2021).

**Fig. 2.**
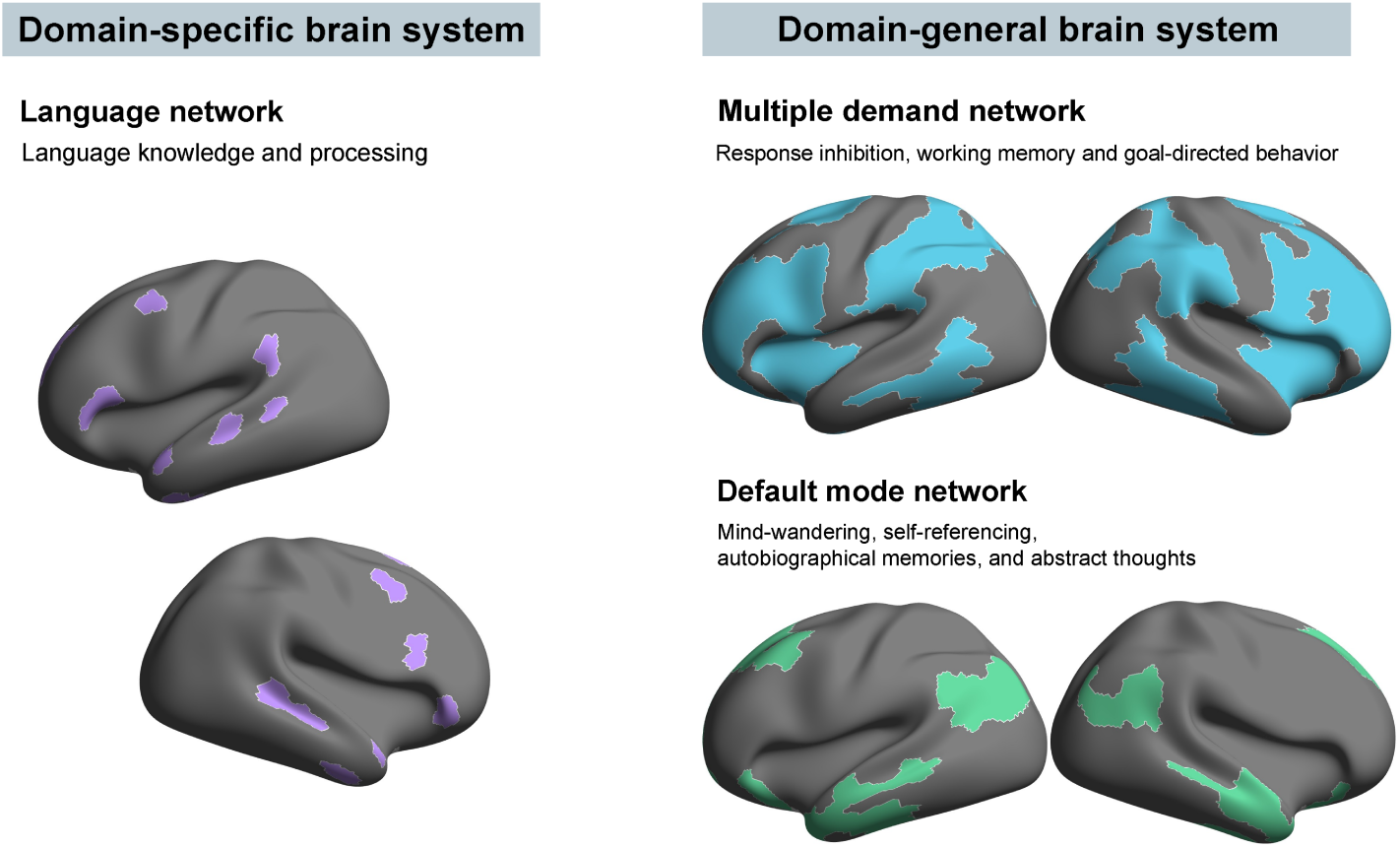
Three Large-scale Brain Systems of Interest. *Note*. The left panel represents the domain-specific language network, while the right panels display domain-general brain system: the MD network (top) and the DM network (bottom).

### Statistical Analyses

Gray matter volumes (GMV) in the three large-scale networks were extracted to test for associations between brain network structure and overall language competence across childhood and adolescence. Data analyses and visualization were performed using SAS 9.4, and packages in R (R Core Team, 2014) respectively.

To investigate developmental associations between cortical morphology and language competence, we employed an exploration-validation framework using two accelerated longitudinal datasets: the devCCNP-PEK Sample as the primary cohort for modeling across childhood and adolescence (6–15 years), and the devCCNP-CKG Sample as an independent validation cohort to enhance result robustness and generalizability through cross-site replication. Developmental trajectory modeling (Section 7) and time-varying effect modeling (Section 7) were conducted solely on the PEK cohort to establish age-related patterns and dynamic associations, leveraging its larger sample for comprehensive analyses. Testing associations in childhood and adolescence (Section 7) applied generalized additive mixed models (GAMMs) to both cohorts, with CKG cohort validating PEK’s findings in the 13–15 years window, a critical period for language and cortical development with robust associations after controlling for non-verbal intelligence, supported by sufficient sample size. The CKG cohort was not used for validation in the 6–8 years window due to insufficient sample size and the less robust associations in the PEK cohort after controlling for non-verbal intelligence, prioritizing robust validation in adolescence to maximize generalizability. Relative Weight Analysis (Section 7) assessed network contributions in both cohorts, with the CKG cohort confirming adolescent results (13–15 years) to ensure consistency across diverse populations.

### Modelling Developmental Trajectories of Language and GMV

To fully model differences in the growth curves of the Verbal Comprehension Index test scores and gray matter volumes on the basis of the precise estimation of curve characteristics, we employed the generalized additive (mixed) model (GAM/GAMM) in our data analyses using the ***mgcv*** (Wood, 2017) package in R (R Core Team, 2014), as this non-parametric regression method does not require a priori knowledge of the associations. GAMM is highly suitable for repeated measurements (Alexander-Bloch et al, 2014; Harezlak et al, 2005), accounting for both within-subject correlation over time and between-subject developmental variations in our accelerated longitudinal design. To more specifically model differences in age trajectories among network-level gray matter volumes and test scores, we employed the following set of GAMMs (see Eq. 1) to detect age-related changes revealed by test scores and gray matter volumes (GMVs) separately:

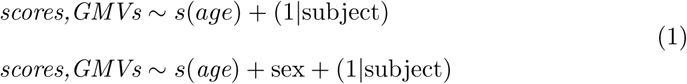

In our models, the smoothing function *s*, employing a fixed degree of freedom and cubic B-splines, was set to have five knots. This number was determined to be optimal for our data, as it achieved a *k* -index close to one without substantial changes in model fit when adjusting the suspected *k* -value (Cho et al, 2024; Wood, 2017). This optimization was guided by the Akaike Information Criterion (AIC). GAMM was fitted with different values of *k* ranging from three to ten, and five was determined to be optimal as it was sufficiently large to represent potential changes from best model fits, and sufficiently small to maintain computational efficiency (Zhou et al, 2021). The first GAMM models gray matter volumes and test scores as smoothing functions of age. Given the evidence of significant sex differences in both intelligence structure and its underlying brain neurophysiological mechanisms (MacDonald et al, 2014), we established the second GAMM with sex as a covariate in the regression. The lowest AIC value was used to determine the best model, and the model-chosen was required to be significantly different from null.

We also performed analyses to test whether there are significant differences in developmental rates among subtest standardized scores and gray matter volumes across the three large-scale neural networks. Specifically, we used partial correlations to account for sex, and conducted a Fisher’s *z* -test (Cohen et al, 1983) using the ***diffcor*** (Cohen, 1988; Steiger, 1980) package in R (R Core Team, 2014). This approach enabled us to investigate potential differences in the pace of language and brain development. Results were evaluated at a significance level of *p < .*05.

### Dynamic Links between the Development of Language and GMV

The Time-Varying Effect Model (TVEM) macro version 3.1.1 (Li et al, 2017; Tan et al, 2012) in SAS 9.4 was used to simulate dynamic associations between network-level gray matter volumes and the Verbal Comprehension Index as age increases, and to locate the temporal window in which the two exhibit a significant correlation. TVEMs were estimated separately for the Verbal Comprehension Index and its three subtests (see Supplementary Materials B for details) and for the three large-scale networks (6 hemispheric measures; see Supplementary Materials F for details). The basic functions of TVEMs were defined using the P-spline penalty for the automatic selection of smoothness and adjustment of independence (Li et al, 2017). One time-variant variable of gray matter volume, and one non-time-varying covariate of sex (0 = Male; 1 = Female) were entered into the model simultaneously to capture temporal effect changes of covariates on the outcome of the Verbal Comprehension Index (or its subtests).

In sum, each TVEM model, run separately for networks and tests, examined the time-varying associations of network-level gray matter volume and overall language competence across ages 6-15 years, controlling for the time-invariant variable of sex. For each network-level gray matter volume, a figure was presented to show how its relationship with the Verbal Comprehension Index changes over time. Pointwise confidence bands created to have 95% confidence using standard techniques were also shown in the figure. Statistically significant associations (*p < .*05) between gray matter volume and the Verbal Comprehension Index were indicated by the criterion (dashed line) falling out of the 95% confidence intervals (CI).

### Linking Language Competence and its Network Basis at Specific Age Groups

Based on TVEM results which determined age-specific subgroups demonstrating statistically significant correlations, we then employed the following set of GAMMs (when the specific age group involved repeated measures) or GAMs (when the specific age group did not involve repeated measures) (see Eq. 2) to more specifically capture the precise estimation of curve characteristics, and to fully model association differences between network-level gray matter volumes (GMVs) and test scores:

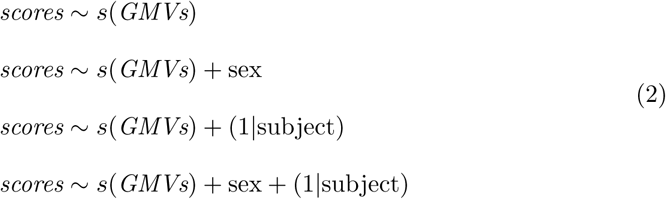

The first GAM was constructed by modeling test scores as a smoothing function of age. The third GAMM incorporated repeated measurements to account for both within-subject dependency and developmental differences among participants. We also established the second GAM and the fourth GAMM by including sex as a fixed-effect term to control for the influence of sex. Given the potential confounding effects of general cognitive abilities on language competence, we also conducted GAMM with non-verbal intelligence as covariate in the regression. Non-verbal intelligence was measured using indices of working memory, processing speed, and perceptual reasoning from the Wechsler Intelligence Scale for Children-IV (WISC-IV), to control for the influence of non-linguistic cognitive abilities on the developmental trajectories of language competence and gray matter volumes. For all of these GAM and GAMM estimations, a threshold of *p <*.05 was used to evaluate significance, and a False Discovery Rate (FDR) correction was applied to correct for multiple comparisons. The CKG Sample was subjected to parallel analyses for verification.

### Measuring Network Contributions to the Development of Language Competence

Additionally, Relative Weight Analysis (RWA) was performed on both samples using the ***rwa*** package in R (R Core Team, 2014) to test the unique and relative contributions of the three large-scale networks in predicting the development of overall language competence throughout childhood and adolescence. RWA transformed correlated predictors into a set of uncorrelated variables maximally relating to the original ones, thereby addressing the problem of multicollinearity (Johnson, 2000; Tonidandel and LeBreton, 2011).

### Exploratory Analyses

To test whether our findings for the MD network generalized across definitions, we repeated the analyses using a task-defined MD map based on hard-versus-easy contrasts from prior work (Fedorenko et al, 2013); see Supplementary Materials C), rather than relying solely on resting-state network parcellations. To assess the specificity of our effects to language-relevant networks, we also repeated analyses using the limbic network from Yeo et al (2011) as a control (Supplementary Materials D). Finally, to contrast our broad measure of language competence with a more narrowly defined experimental task, we compared VCI scores with performance on a lab-based audiovisual integration task (Supplementary Materials E). These exploratory analyses were limited to the PEK sample and served as supplementary tests of robustness, rather than central components of the study.

## Results

### Developmental Trajectories of Language and GMV

The best-fitting age trajectory for the Verbal Comprehension Index determined by AIC in the PEK Sample included sex as a covariate; this was significantly different from the null model for the age effect (see Table 2). GAMM estimates indicated an improvement in language ability with increasing age, see Figure 3a.

**Fig. 3.**
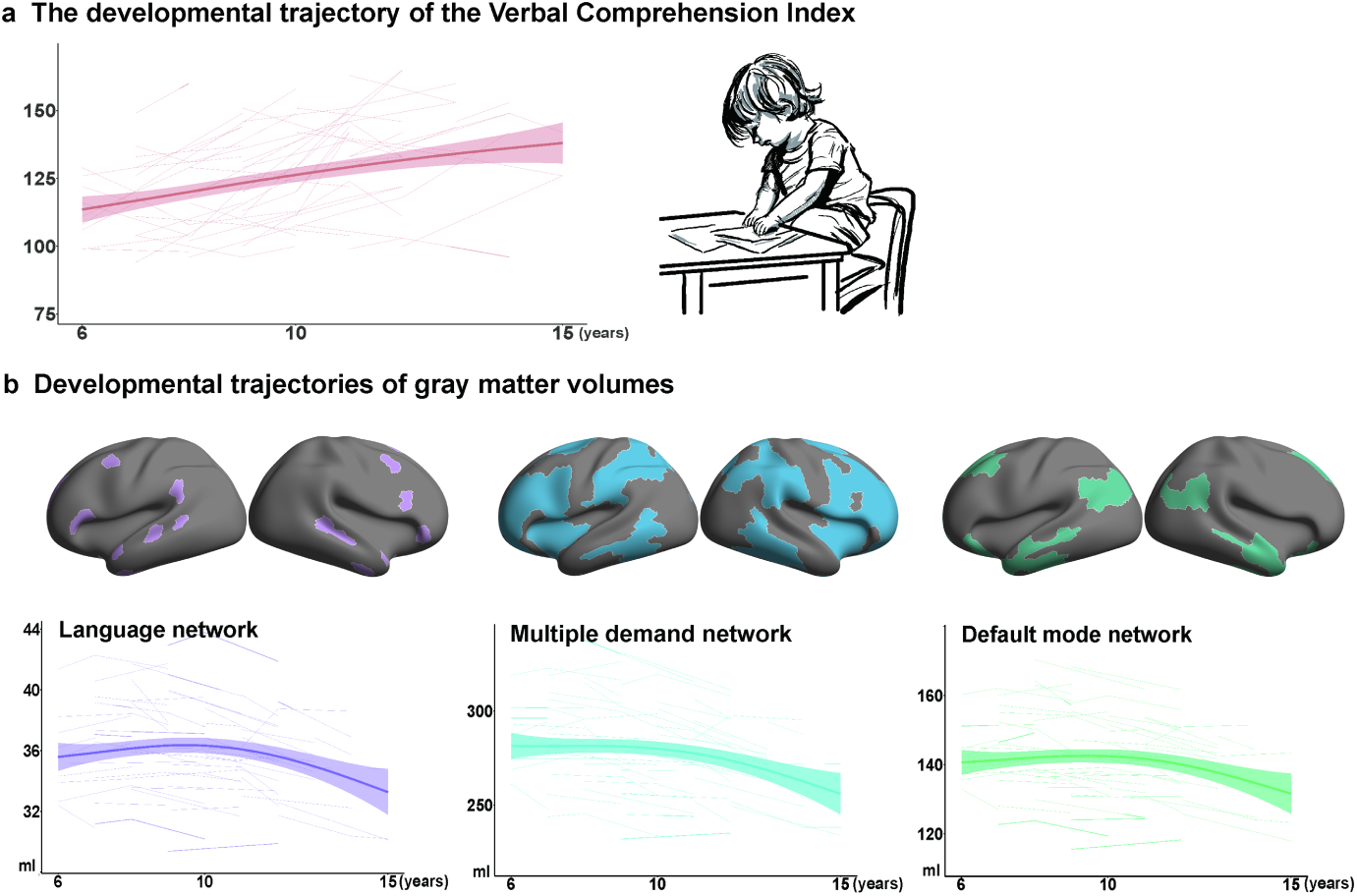
Developmental Trajectories of Language and GMV. *Note*. **a**: The developmental trajectory of the Verbal Comprehension Index (red). **b**: Developmental trajectories of gray matter volumes across three large-scale networks. Purple represents the trajectory of volumetric changes in the language network, blue signifies the trajectory of the multiple demand network, and green denotes the trajectory of the default mode network.

**Table 2.**
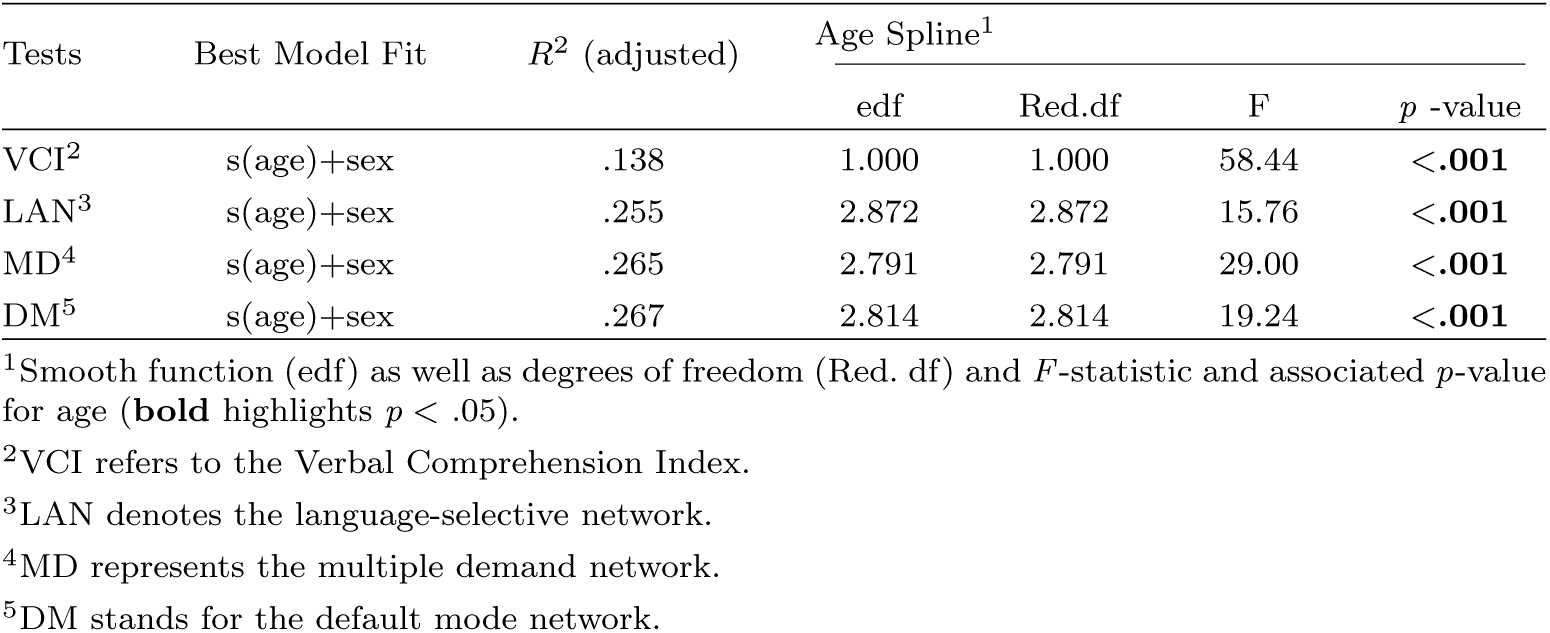
GAMM Estimates for the Verbal Comprehension Index and GMV for the PEK Sample.

| Tests | Best Model Fit | $R^2$ (adjusted) | Age Spline<sup>1</sup> | | | |
| --- | --- | --- | --- | --- | --- | --- |
| | | | edf | Red.df | F | $p$ -value |
| VCI<sup>2</sup> | s(age)+sex | .138 | 1.000 | 1.000 | 58.44 | <.001 |
| LAN<sup>3</sup> | s(age)+sex | .255 | 2.872 | 2.872 | 15.76 | <.001 |
| MD<sup>4</sup> | s(age)+sex | .265 | 2.791 | 2.791 | 29.00 | <.001 |
| DM<sup>5</sup> | s(age)+sex | .267 | 2.814 | 2.814 | 19.24 | <.001 |

<sup>1</sup>Smooth function (edf) as well as degrees of freedom (Red. df) and $F$ -statistic and associated $p$ -value for age (**bold** highlights $p < .05$ ).
<sup>2</sup>VCI refers to the Verbal Comprehension Index.
<sup>3</sup>LAN denotes the language-selective network.
<sup>4</sup>MD represents the multiple demand network.
<sup>5</sup>DM stands for the default mode network.

For network-level gray matter volumes, AIC indicated that the best-fitting model was the one including the sex term (see Table 2). GAMM-based estimates revealed a significant non-linear decreasing trend for the development of gray matter volume across three large-scale networks (language: *p <* .001; MD: *p <* .001; DM: *p <* .001, see Figure 3 and Table 2). This non-linear trend indicates a complex maturation process, characterized by an initial increase in volume followed by a pronounced reduction, reflecting the brain’s dynamic restructuring during critical developmental phases. We also conducted Fisher’s *z* -test in a post-hoc step to reveal differences in age-related development among the three networks. The MD network demonstrated a more rapid decline in gray matter volume (*r* = -.240) across childhood and adolescence than the other two networks (language network: *Z* = 4.02, *p <*.001; DM network: *Z* = 7.207, *p <* .001). Post-hoc volume-wise GAMMs further indicated that both the left and right hemispheric volumes obtained separately from the three networks showed similar nonlinear developmental patterns (see Supplementary Materials F).

### The Development of Language Competence and its Morphological Basis

#### Dynamic Links between the Development of Language and GMV

The TVEM analysis for the PEK Sample revealed significant positive associations between gray matter volumes in the three large-scale networks and the Verbal Comprehension Index in the younger age group (ages between six and eight years; the language network: 6-7.54 years, the MD network: 6-7.72 years, the DM network: 6-7.72 years), as shown in Figure 4. There were also significant positive relationships between gray matter volumes in the two domain-general networks and the Verbal Comprehension Index in the older age group (age between 13 and 15 years; the MD network: 12.9-15 years, the DM network: 13.09-14.54 years).

**Fig. 4.**
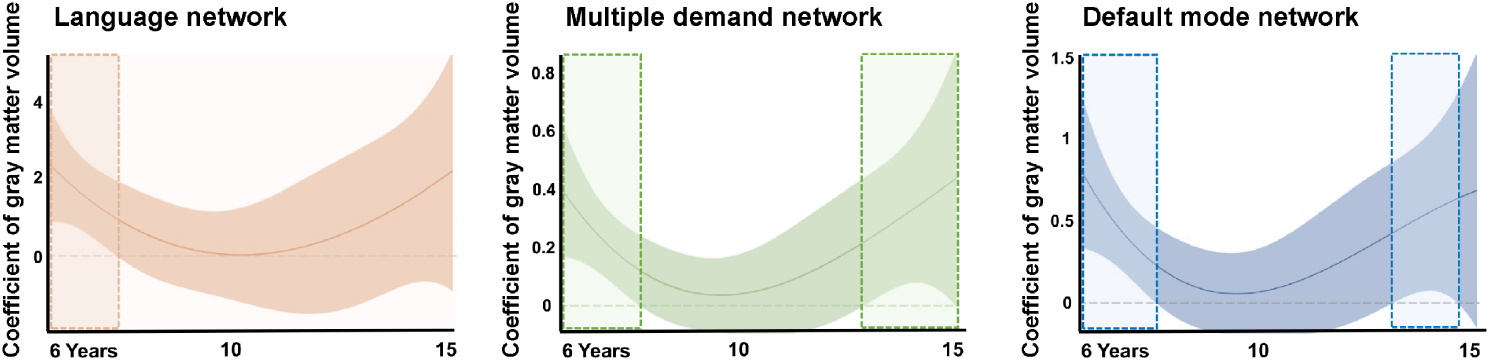
Age-related Associations between GMV and the Verbal Comprehension Index. *Note*. TVEM analysis of age-related associations between the Verbal Comprehension Index and volumetric changes in the language network (red), the multiple demand network (green), and the default mode network (blue) in the PEK Sample. Shaded areas indicate upper and lower 95% confidence intervals. Statistically significant associations (*p < .*05) between gray matter volumes and the Verbal Comprehension Index are indicated by the dotted criterion line falling out of the 95% confidence intervals. Dashed boxes highlight ages at which significant positive associations were observed between gray matter volume and the Verbal Comprehension Index (the 95% confidence band is above the estimated association of 0 on the *y* -axis).

#### Language Competence and its Network Basis at Specific Age Groups

Next, GAMs were applied to the PEK Sample to illustrate the positive associations between gray matter volumes in the three functional networks and the Verbal Comprehension Index in the younger children and adolescents separately. All best-fitting models were determined to be those that controlled for sex based on the AIC (see Table 3). GAM estimates revealed that both the younger children and adolescents exhibited linear positive associations (see Figure 5a). Gray matter volumes in both the domain-specific language network and domain-general MD network and DM network showed significant positive associations with the Verbal Comprehension Index in younger children (the language network: *p <* .05; the MD network: *p <* .05; the DM network: *p <* .05, Table 3), revealing that greater gray matter volume in all three networks is linked to enhanced language skills in children. However, these effects were no longer significant after controlling for non-verbal intelligence indices (measured by the WISC) including working memory, perceptual reasoning, and processing speed (the language network: *p* = .18; the MD network: *p* = .08; the DM network: *p* = .14). Between ages 13 and 15, gray matter volumes in domain-general networks showed robust positive associations with the Verbal Comprehension Index (the MD network: *p <* .05; the DM network: *p <* .05) For GAM estimates, see Table 3; detailed information regarding hemispheric results is provided in Supplementary Material F. These associations remained significant after controlling for non-verbal intelligence (the MD network: *p <* .01; the DM network: *p <* .05).

**Fig. 5.**
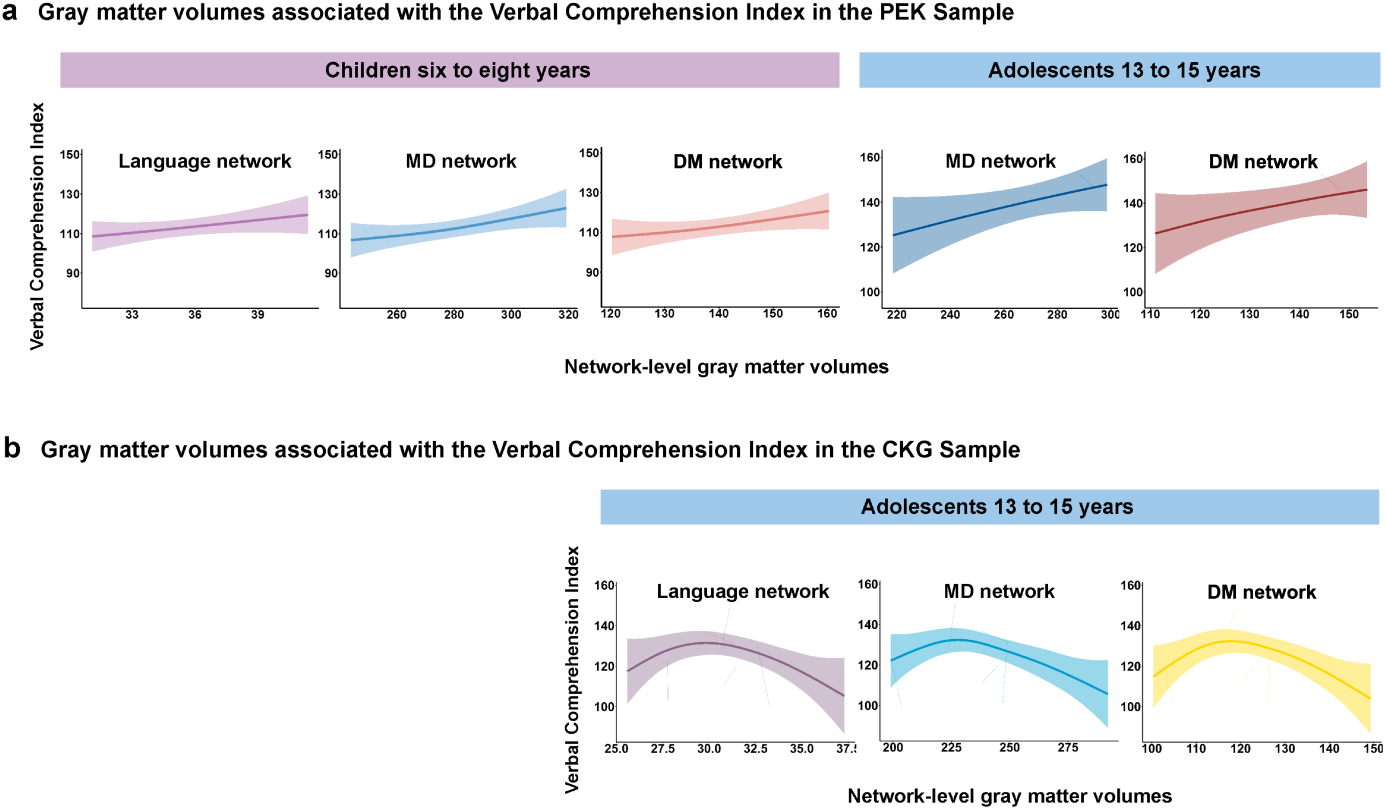
Associations between Language Competence and its Network Basis at Specific Age Groups. *Note*. **a**: Gray matter volumes (the *x* -axis) associated with the Verbal Comprehension Index (the *y* -axis) in specific age groups in the PEK Sample. Panels on the left present results in younger children, while those on the right focus on adolescents. **b**: Gray matter volumes (the *x* -axis) associated with the Verbal Comprehension Index (the *y* -axis) during adolescence in the CKG Sample.

**Table 3.** GAM/GAMM Estimates for Language Competence and its Network Basis at Specific Age Groups.

| Group | Network | Best Model Fit | $R^2$ (adjusted) | Slope<sup>1</sup> | | | |
| --- | --- | --- | --- | --- | --- | --- | --- |
| | | | | edf | Red.df | F | $p$-value |
| The PEK Sample |  |  |  |  |  |  |  |
| Age 6-8 | LAN<sup>2</sup> | s(GMV<sup>5</sup>)+sex | .057 | 1.000 | 1.000 | 4.15 | <.05(.18)<sup>6</sup> |
|  | MD<sup>3</sup> | s(GMV)+sex | .157 | 1.314 | 1.562 | 6.28 | <.05(.08) |
|  | DM<sup>4</sup> | s(GMV)+sex | .089 | 1.002 | 1.003 | 5.49 | <.05(.14) |
| Age 13-15 | MD | s(GMV)+sex | .303 | 1.000 | 1.000 | 9.337 | <.05(<.01) |
|  | DM | s(GMV)+sex | .273 | 1.002 | 1.003 | 8.338 | <.05(<.05) |
| The CKG Sample |  |  |  |  |  |  |  |
| Age 13-15 | LAN | s(GMV)+sex | .166 | 2.674 | 3.175 | 2.96 | .04(.07) |
|  | MD | s(GMV)+sex | .168 | 2.626 | 3.102 | 3.16 | .04(.05) |
|  | DM | s(GMV)+sex | .183 | 2.717 | 3.217 | 3.37 | .04(.05) |

<sup>1</sup>Smooth function (edf) as well as degrees of freedom (Red. df) and $F$ -statistic and associated $p$ -value for gray matter volumes (**bold** highlights $p < .05$ ).
<sup>2</sup>LAN: the language network.
<sup>3</sup>MD: the multiple demand network.
<sup>4</sup>DM: the default mode network.
<sup>5</sup>GMV indicates network-level gray matter volumes.
<sup>6</sup>For the column of $p$ -value in the table, the values in parentheses represent $p$ -values after controlling for non-verbal intelligence.

Analyses of the PEK data indicate that the results for adolescence are the most robust, remaining significant even after controlling for non-verbal intelligence. To further validate the robustness of these findings, we examined the same age window using the CKG dataset. GAMM analyses demonstrated significant associations between the Verbal Comprehension Index and gray matter volumes in all three networks during early adolescence (the language network: *p* = .04; the MD network: *p* = .04; the DM network: *p* = .04), as shown in Figure 5b and Table 3. After controlling for non-verbal intelligence, the associations between the Verbal Comprehension Index and gray matter volumes in the MD network (*p* = .05) and the DM network (*p* = .05) remained significant, while the association with the language network was not significant (*p* = .07). This implies that, during early adolescence, further enhancement in the ability to use language for reasoning and expression is more closely related to gray matter volumes in domain-general networks. However, the form of this relationship differed: while the PEK data showed a linear positive association, the CKG cohort exhibited a non-linear (inverted U-shaped) relationship, with the highest language scores associated with intermediate levels of gray matter volume. This suggests that the contribution of domain-general networks to language competence may not scale linearly with volume across all individuals or populations, potentially following the non-linear developmental trajectory of verbal intelligence observed in the CKG Sample (see Figure S2 for details).

#### Network Contributions to the Development of Language Competence

RWA was conducted for both samples to estimate the unique contributions of each network to the enhancement of overall language competence resulting from volumetric variations (Table S2). Results indicated that the development of gray matter volume within the MD network was the most important volumetric factor facilitating overall language development, as evidenced by its high relative weights in the PEK Sample (relative weights in children aged six to eight: .11; relative weights in adolescents aged 13 to 15: .13) and the CKG Sample (relative weights in adolescents aged 13 to 15: .07). Specifically, the MD network accounted for 46% of the increase in the Verbal Comprehension Index caused by volumetric changes in children. During adolescence, it also accounted for 57% of the change in Verbal Comprehension in the PEK Sample and 48% in the CKG Sample, see Figure 6.

**Fig. 6.**
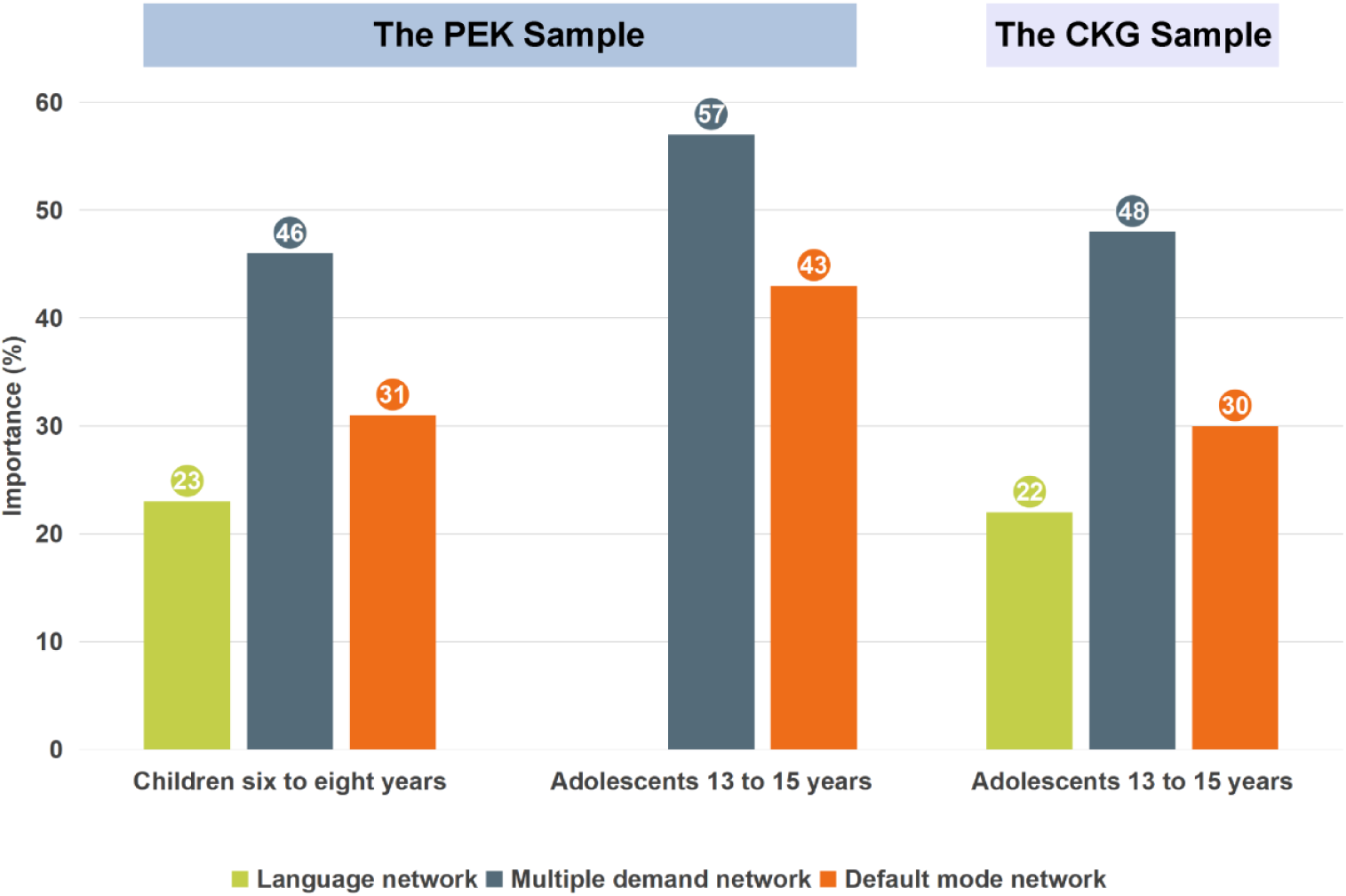
Relative Weights Analysis of Network GMV Predicting Language Competence. *Note*. Importance (%): rescaled relative weights (i.e., relative weights divided by full model *R*^2^). We included here the networks whose volumes were significantly associated with the Verbal Comprehension Index in our analysis of relative weights. For the PEK Sample, we calculated the relative weights of all the three large-scale networks during childhood and concentrated on domain-general networks during adolescence, with their combined total equaling 100%. For the CKG Sample, we computed the relative weight of all the three network for adolescents.

### Language Competence and its Network Basis at the Subtest Level

Analysis of changes in gray matter volume associated with the three subtests revealed results for the Vocabulary subtest that paralleled the Verbal Comprehension Index in younger children and adolescents, with significant positive volumetric correlations across childhood and adolescence. For the Similarities subtest, we found significant positive associations in children aged approximately six to ten across all the three large-scale networks. However, no significant volumetric correlates were found for the Similarities subtest during adolescence. The Comprehension subtest showed no significant age-related interactions with volumetric development at the network level. Detailed information about the subtests and their volumetric correlates can be found in Supplementary Material B. Post-hoc analyses revealing hemispheric association patterns are presented in Supplementary Material F.

Regarding network contributions, the gray matter volume in the language network emerged as the most important factor across childhood. It explained more than 40% of the enhancement in the subtest scores (Vocabulary: 43.7%; Similarities: 43.9%) attributable to variations in network-level gray matter volumes. In adolescents, changes in the MD network were the primary predictor of enhancements in Vocabulary scores, explaining more than 54.5% of the variance in the outcomes. Additionally, the gray matter volume in the DM network contributed significantly, on average accounting for 35.0% of the enhancement of subtest scores during the two developmental periods. For further details, please refer to Supplementary Materials B.

### Results for Exploratory Analyses

#### Results for the Evelab MD System

Parallel analyses were performed using gray matter volumes extracted from the “Evelab MD system”. The results successfully replicated our main findings regarding dynamic associations between gray matter volumes in the MD system during two developmental periods (childhood aged six to eight and adolescence aged twelve to fifteen) and the significant contribution of this predefined MD system to the development of overall language competence (see Supplementary Materials C for details). These findings confirm the methodological validity of the MD network used in the present study.

#### Results with the Limbic Network as a Control System

TVEM analysis with the limbic system as a control network revealed relatively weak correlation between gray matter volume and the Verbal Comprehension Index. GAM estimates revealed no significant association between the development of language performance and the development of gray matter volume in the limbic system. These results highlight the specificity of our main results (Supplementary Materials D).

#### Results for the Audiovisual Integration Task

GAM estimates revealed marginally significant correlations between performance in the audiovisual integration task and both the language network and the MD network during childhood, with the language network exhibiting substantially greater weights compared to the multiple-demand network. During adolescence, the language network emerged as the only significant predictor of audiovisual integration before correction. These results demonstrate that the language network significantly contributes to core language functions. The strong contributions of the multiple demand network and the default mode network reported above are likely attributable to our use of verbal intelligence as the measure of language ability linked to higher-order thought (Supplementary Materials E).

## Discussion

This study addresses a long-standing debate about whether language competence depends primarily on domain-specific mechanisms or is shaped by broader cognitive systems. Taking a developmental perspective, we provide empirical support for the domain-general view by showing that structural variation in domain-general networks – particularly the default mode and multiple-demand systems – plays a key role in the emergence of verbal reasoning and expression skills. Using two large-scale longitudinal cohorts, we found that language competence develops through the joint contribution of domain-specific and domain-general networks, with domain-general systems exerting a stronger influence during key windows of adolescence. These effects were not uniformly distributed across development but instead emerged during specific periods when maturation of large-scale cortical networks aligned with gains in verbal ability. This developmental coupling between structural brain maturation and language competence highlights the importance of general-purpose cognitive systems in supporting language use in real-world contexts. Our findings converge with prior evidence that complex language behavior draws on domain-general systems, especially in tasks that extend beyond passive comprehension (Fedorenko et al, 2024a; Fedorenko and Varley, 2016; Mahowald et al, 2024).

The strong contribution of domain-general networks to language competence across both cohorts likely reflects two key factors: the nature of the verbal tasks and the developmental stage of the participants in our study. The Verbal Comprehension Index includes subtests that go beyond basic linguistic decoding to tap verbal reasoning, conceptual knowledge, and real-world semantic understanding (Lynne Beal, 2004; Wechsler, 2003). Verbal concept information and verbal reasoning relate to the way that language is deployed in everyday situations, consistent with the role of general-purpose resources to answer questions (e.g., Diachek et al, 2020; Fedorenko et al, 2024a; Hu et al, 2023). These broader demands are consistent with the observed involvement of the MD and DMN networks, which support flexible, goal-directed cognition and semantic integration. Children and adolescents might rely more heavily on these general-purpose systems for language processing than adults, whose neural architecture may afford more efficient or specialized language pathways. This interpretation aligns with previous evidence that language networks in youth are less functionally specialized than in adulthood, showing more diffuse activation (Brauer et al, 2010; Weiss et al, 2018) and greater power across broad EEG bands (Schneider et al, 2018). In line with these findings, a systematic review (Weiss-Croft and Baldeweg, 2015) and developmental studies (Rosselli et al, 2014) have highlighted ongoing interactions between language development and non-linguistic functions including executive control, memory, and social cognition. Taken together, these results support the view that in earlier stages of development, language competence in demanding contexts depends not only on language-selective regions but also on the broader scaffolding of domain-general cognitive systems (Fedorenko et al, 2024a).

Our findings also show that gray matter volume predicted overall language competence only during specific developmental windows, rather than consistently across the full span of childhood and adolescence, as revealed by TVEM. These findings are consistent with prior research indicating major changes in intelligence (Shaw et al, 2006) and language functions (Blakemore and Mills, 2014; Budisavljevic et al, 2015; Hattouti et al, 2019; Lidzba et al, 2011; Nippold, 2000; Ricketts et al, 2020; Smith, 2015; Wassenberg et al, 2008) that originate in childhood and extend well into adolescence. Moreover, the identification of significant contributions of gray matter volume to the development of overall language skills at certain developmental stages highlights the complex interplay between brain maturation and language development in both children (Fengler et al, 2015) and adolescents (Budisavljevic et al, 2015). This complexity is highlighted by differences between the cohorts: although they both demonstrated that domain-general networks are linked to language competence during adolescence, the nature of this relationship differed. In the Beijing cohort, greater gray matter volume in the MD and DM networks was linearly associated with better performance on the Verbal Comprehension Index, replicating established association patterns (e.g., Torre and Eden, 2019). In contrast, the Chongqing cohort exhibited a nonlinear pattern. We propose this divergence may reflect neurodevelopmental adaptations associated with population-specific dual-language acquisition, as the concurrent learning of Mandarin Chinese and Chongqing dialect appears to drive the unique developmental trajectory of verbal intelligence in this population. The above suggests that the relationship between structure and function is dynamic, with the effectiveness of domain-general systems in supporting language depending on both their maturity and the cognitive demands of the task across individuals and populations.

Our study highlights how linguistic and cognitive task demands impact the contributions made by the domain-specific language network and domain-general cognitive networks. Through subtest analyses, we found that the Vocabulary and Similarities subtests, which effectively predict core linguistic aspects like lexicon and word knowledge, have the language network as their most crucial predictor. In contrast, the Comprehension subtest, which serves as a more sophisticated predictor of higher-level language skills involving practical knowledge and judgement based on past experiences (Weiss et al, 2015), relies more on domain-general factors. Moreover, in contrast to the prominence of general-purpose brain networks in our main analyses, an audiovisual integration task identified the language network as the most significant contributor to performance. This aligns with previous studies showing functional specificity of the language-selective network in tasks that focus on a specific linguistic domain (Fedorenko et al, 2011; Fedorenko and Shain, 2021; Mineroff et al, 2018). While domain-specific regions may suffice for core linguistic computations (e.g., syntactic parsing; Blank and Fedorenko, 2017), real-world language competence—particularly during development—relies on the integration of domain-general resources to support goal-directed communication (Fedorenko et al, 2024b). This is consistent with the notion that language serves as a tool for thought (Fedorenko and Varley, 2016), with its neural implementation flexibly engaging either specialized or general-purpose systems depending on task demands and developmental stage. Experimental tasks designed to isolate linguistic processes might not capture the complexities of language, since tight control removes crucial aspects of language use from tasks. These findings highlight the importance of considering more naturalistic language use when investigating the neural basis of language processing.

Despite the close link between overall language development and domain-general volumetric correlates identified in some developmental periods, further research is needed to establish the extent to which our findings would generalize to tasks with different demands, and whether the role of the multiple demand and the default mode networks reflects the executively-challenging nature of verbal comprehension during childhood and adolescence. Incorporating a broader range of measures, including white matter connectivity (Qi et al, 2019) and connectivity gradients (Dong et al, 2021, 2024), would provide a more comprehensive picture of the relationship between network-level brain development and real-life language performance.

## Conclusion

Taken together, our findings support the domain-general view by demonstrating that language competence relies on both language-selective and domain-general networks, with the latter playing a more prominent role during specific developmental periods. This challenges the domain-specific perspective that language is solely subserved by specialized neural circuits. Importantly, these effects were observed alongside the progressive development of language performance (Berken et al, 2017; Nayak et al, 2022; Rosselli et al, 2014), highlighting the dynamic interplay between brain maturation and language acquisition. Our study emphasizes the need to consider domain-general cognitive resources in models of language development, particularly during sensitive periods of neuroplasticity.

## Acknowledgement

This research is supported by the STI 2030—the major projects of the Brain Science and Brain-Inspired Intelligence Technology (2021ZD0200500), the Natural National Science Foundation of China (Grant/Award Numbers: 31871108), open Research Fund of the State Key Laboratory of Cognitive Neuroscience and Learning, the Fundamental Research Funds for the Central Universities, the Research Funds of Renmin University of China (23XNKJ01).

