## supplemental files for "Domain-General Brain Networks Support Language Development"

### Supplementary Materials

#### A Sample Distributions and Verbal Intelligence Development across Cohorts

##### A1 Sample Distributions in the PEK and CKG Samples

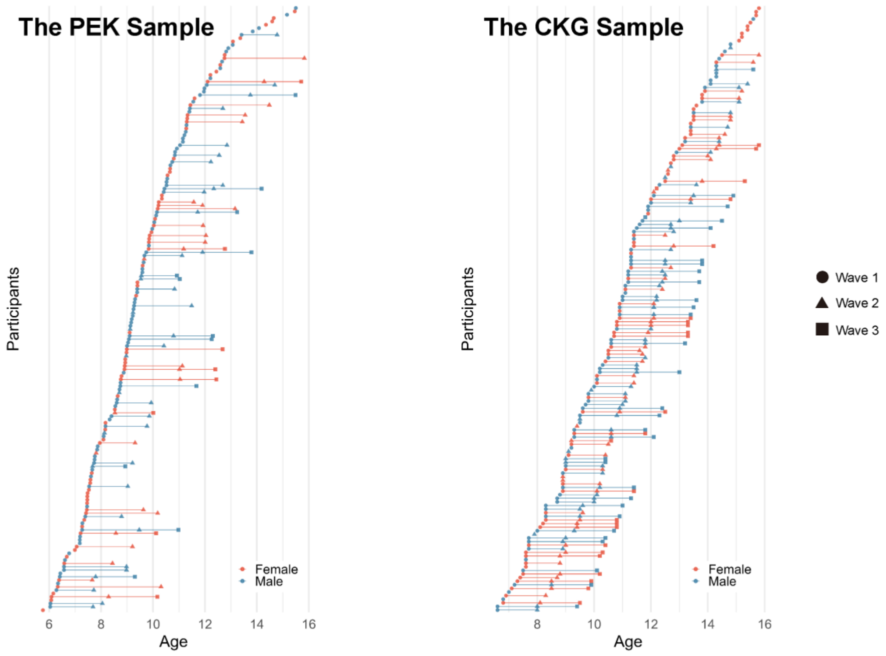


**Fig. S1 Age and sex Distributions in the PEK and CKG Samples**.

*Note*. The left panel displays age and sex distributions for participants' completion in the PEK Sample, while the right panel shows age and sex distributions in the CKG Sample. Females are represented in red and males in blue. Circular markers indicate data from the first measurement wave, triangles denote the second wave, and squares represent the third wave of data collection.

#### A2 Age-related Verbal Intelligence Development in the PEK and CKG Samples

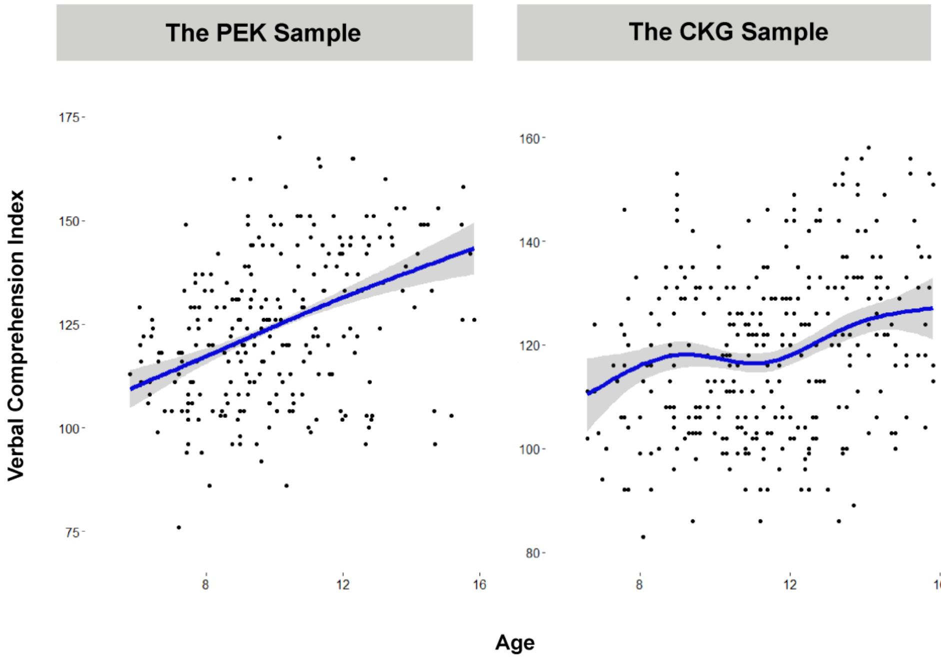


**Fig. S2 Developmental Trajectories of Language across Cohorts.**

*Note.* The left panel presents age-related developmental trajectory of the Verbal Comprehension Index in the PEK Sample, while the right panel presents the corresponding developmental patterns observed in CKG Sample.

### B Results of GMV Development and the Three Subtests

#### B1 Developmental Trajectories of the Three Subtests

The growth patterns for the Vocabulary and Comprehension subtests showed a significant increase across childhood and early adolescence, implying a continuous improvement in language skills. In contrast, the growth curve for the Similarities subtest displayed a slightly inverted U shape, showing a minor decline during early adolescence (ages 12 to 15 years), as shown in Figure S3. However, this decline was not significant statistically when tested separately ($p = .155$). Additionally, a Fisher's $z$-test was conducted to identify significant differences in age-related development between the Comprehension subtest and the other two subtests (Comprehension vs. Vocabulary: $Z$ = -2.024, $p$ = .043; Comprehension vs. Similarities: $Z$ = -2.865, $p$ = .004). Notably, the Comprehension subtest demonstrated a more substantial increase ($r$ = .358) throughout childhood and adolescence.

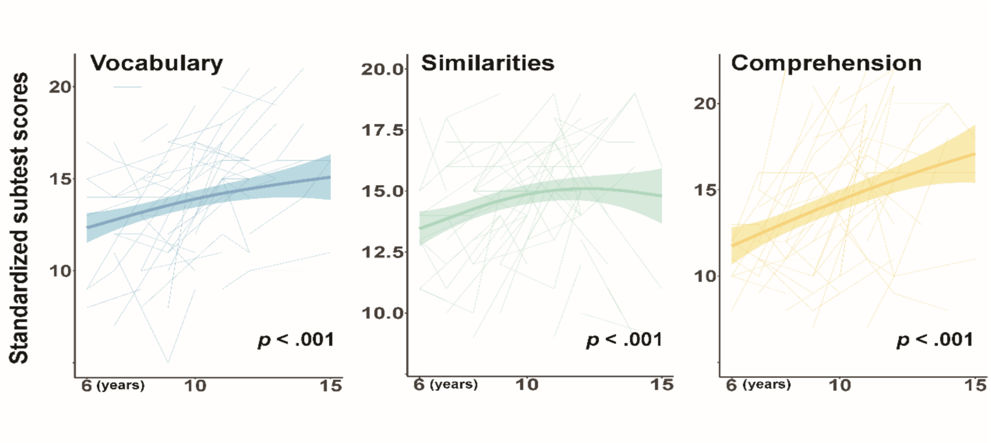


**Fig. S3 Developmental Trajectories of the Three Subtests in the Full Sample.**

*Note.* The blue color represents the trajectory for the Vocabulary subtest, the green color denotes the trajectory for the Similarities subtest, and the yellow color indicates the trajectory for the Comprehension subtest in the full sample.

#### B2 Correlation between GMV Changes and the Three Subtests

TVEM analysis of the Vocabulary subtest revealed an association pattern similar to that of the Verbal Comprehension Index. Significant positive associations were identified between the Vocabulary subtest score and gray matter volumes in all the three networks among early school-age children (ages 6-7), and between the score and two domain-general networks in adolescents (ages 13-15). For the Similarities subtest, significant positive associations were observed in children aged approximately six to ten years across the three networks. Additionally, the Comprehension subtest showed no significant age-related interactions with gray matter volume development, see Figure S4a.

GAM estimates indicated that in the youngest subgroup, the Vocabulary subtest score increased significantly with gray matter volume development across all three networks. In adolescents aged 13 to 15, the association curves in the two domain-general cognitive networks were flatter but still reached statistical significance after correction (see Figure S4b and Table S1), suggesting a continued role of these general-purpose resources in vocabulary development. For the Similarities subtest, GAMM analyses revealed significant linear positive associations across all three networks during the school years (see Figure S4c and Table S1), indicating that superior Similarities abilities are linked to increased gray matter volumes in these networks. Post-hoc analysis revealed similar association patterns in hemispheric gray matter volumes, implying the stability of the effect (see supplementary material F).

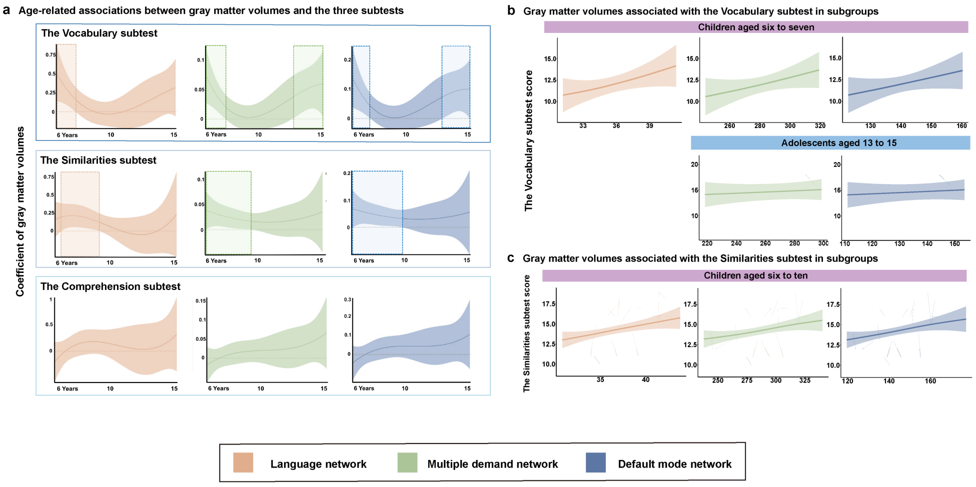


**Fig. S4 Age-related Associations between GMV and the Three Subtests.**

*Note.* **a**: Age-related associations between the three subtest scores and volumetric changes in the language network (red), the multiple demand network (green), and the default mode network (blue). Shaded areas indicate upper and lower 95% confidence intervals. Statistically significant associations ($p < .05$) between gray matter volumes and subtest scores are indicated by the criterion dotted line falling out of the 95% confidence intervals. Dashed boxes highlight ages at which significant positive associations were observed between gray matter volume and subtest scores (as indicated by the fact that the 95% confidence band is above the estimated association of 0 on the $y$-axis). The top panel presents results for the Vocabulary subtest, the middle panel for the Similarities subtest, and the bottom panel for the Comprehension subtest. **b**: Gray matter volumes associated with the Vocabulary subtest score across childhood (top) and adolescence (bottom). The red color indicates the association with the language network, green represents the multiple demand network, and blue denotes the default mode network. **c**: Gray matter volumes associated with the Similarities subtest score in children aged six to ten. The red color indicates the association with the language network, green represents the MD network, and blue denotes the DM network.

**Table S1**

GAM/GAMM Estimates for GMV Associated with the Three Subtests in Subgroups.

| | Group | Network | $R^2$ (adjusted) | Slope[1] | | | |
|---|---|---|---|---|---|---|---|
| | | | | **edf** | **Red.df** | **F** | ***p*-value** |
| Voc[2] | Age 6-7 | LAN[4] | .2350 | 1.198 | 1.368 | 5.595 | <**.05** |
| | | MD[5] | .2190 | 1.379 | 1.665 | 5.501 | <**.05** |
| | | DM[6] | .1940 | 1.000 | 1.000 | 7.010 | <**.05** |
| | Age 13-15 | MD | .2150 | 1.000 | 1.000 | 5.360 | <**.05** |
| | | DM | .2230 | 1.000 | 1.000 | 5.640 | <**.05** |
| Sim[3] | Age 6-10 | LAN | .0741 | 1.000 | 1.000 | 8.164 | <**.05** |
| | | MD | .0539 | 1.000 | 1.000 | 6.018 | <**.05** |
| | | DM | .0577 | 1.000 | 1.000 | 6.339 | <**.05** |

[1]Smooth function (edf) as well as degrees of freedom (Red. df) and $F$-statistic and associated $p$-value for gray matter volume (**bold** highlights $p < .05$).

[2]Voc stands for the Vocabulary subtest.

[3]Sim refers to the Similarities subtest.

[4]LAN refers to the language-selective network.

[5]MD denotes the multiple demand network.

[6]DM represents the default mode network.

#### B3 Network Contributions to Specific Linguistic Aspects

For the two subtests, RWA revealed that the gray matter volume in the language network was the most important contributing factor across childhood. It explained more than 40% of the enhancement in subtest scores (the Vocabulary subtest: 43.71%; the Similarities subtest: 43.85%) resulting from variations in network-level gray matter volume. Nevertheless, changes in the MD network were still the most important factor in predicting enhancement in Vocabulary score in adolescents, explaining more than 54.53% of the variance in the outcomes. Moreover, gray matter volume in the DM network also played an important role in the two subtests in both the two critical periods. Detailed information is presented in Figure S5 and Table S2.

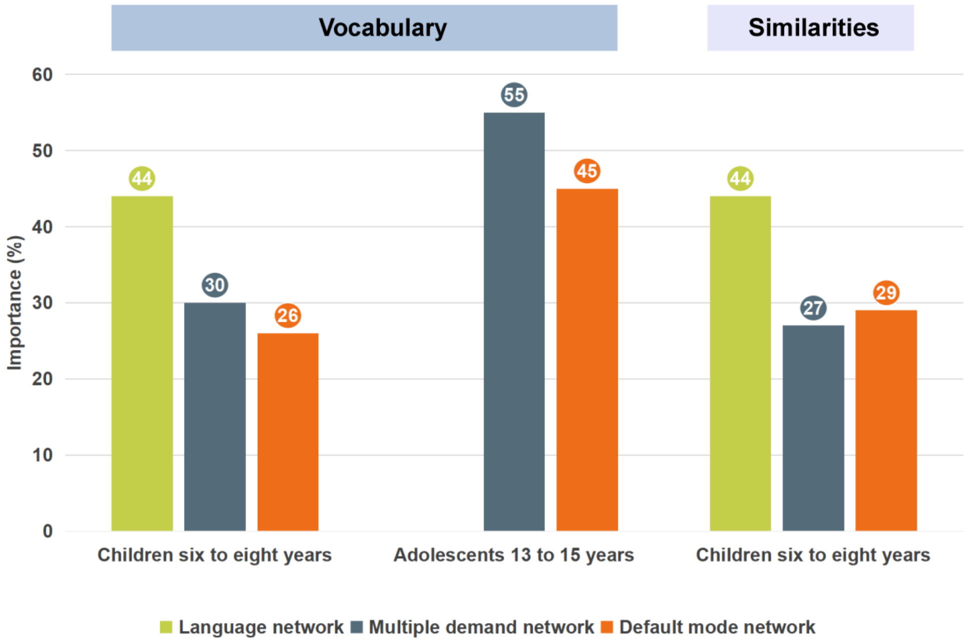


**Fig. S5 Network GMV Contributions to Specific Linguistic Aspects**.

*Note*. Importance (%): rescaled relative weights (i.e., relative weights divided by full model $R^2$). We included here the networks whose volumes were significantly associated with the subtest scores in our analysis of relative weights. For children aged six to eight, we calculated the relative weights of all the three large-scale networks. For adolescents, we concentrated on the relative weights of the MD network and the DM network, with their combined total equaling 100%.

**Table S2**

Relative Weight Analysis of Network GMV Predicting Language Development

| Network | Group | Relative Weights[1] | Importance (%)[2] | $\delta$ | Rank |
|---|---|---|---|---|---|
| **The PEK Sample** | | | | | |
| Dependent Variable: the Verbal Comprehension Index | | | | | |
| LAN[3] | | .055 | 23.09 | .091 | 3 |
| MD[4] | Age 6 - 8 | .110 | **46.06**[6] | .478 | 1 |
| DM[5] | | .073 | 30.85 | .028 | 2 |
| $R^2 = .24$ | | | | | |
| MD | Age 13 - 15 | .129 | **56.57** | .434 | 1 |
| DM | | .091 | 43.43 | .202 | 2 |
| $R^2 = .23$ | | | | | |
| Dependent Variable: the Vocabulary Subtest Score | | | | | |
| LAN | | .076 | **43.71** | .325 | 1 |
| MD | Age 6 - 7 | .053 | 30.37 | .235 | 2 |
| DM | | .045 | 25.92 | .111 | 3 |
| $R^2 = .17$ | | | | | |
| MD | Age 13 - 15 | .077 | **54.53** | .318 | 1 |
| DM | | .064 | 45.47 | .201 | 2 |
| $R^2 = .14$ | | | | | |
| Dependent Variable: the Similarities Subtest Score | | | | | |
| LAN | | .042 | **43.85** | .250 | 1 |
| MD | Age 6 - 10 | .026 | 27.06 | .106 | 3 |
| DM | | .028 | 29.10 | .147 | 2 |
| $R^2 = .10$ | | | | | |
| **The CKG Sample** | | | | | |
| Dependent Variable: the Verbal Comprehension Index | | | | | |
| LAN | | .030 | 21.71 | -.054 | 3 |
| MD | Age 13 - 15 | .066 | **47.66** | -.362 | 1 |
| DM | | .043 | 30.63 | .072 | 2 |
| $R^2 = .14$ | | | | | |

[1]Relative Weights is an estimated $R^2$ associated with each volumetric predictor.

[2]Importance (%): rescaled relative weights (relative weights divided by full model $R^2$).

[3]LAN: the language-selective network.

[4]MD: the multiple demand network.

[5]DM: the default mode network.

[6]**Bold** indicates the strongest estimated predictor.

#### C Results Obtained with the MD Regions Defined by Fedorenko et al (2013)

Using the group-level representation of brain regions throughout the MD system from average activity reported by Fedorenko et al (2013), we asked if our findings regarding the MD network incorporated in the main text replicate, defined on the basis of co-activation of seven brain networks (Salience/VenAttn A; Salience/VenAttn B; Control A; Control B; Control C; Dorsal Attention A; Dorsal Attention B) from Kong et al (2021). The predefined multiple brain regions in the MD system, which we referred to as the "Evelab MD system" were presented in Figure S6a (Fedorenko et al, 2013), (https://imaging.mrc-cbu.cam.ac.uk/imaging/MDsystem).

Using the same analytical methods covering TVEM and GAMM/GAM, we replicated our main findings reported in the main text with this predefined MD system. GAMM-based estimates revealed that the developmental trajectory of the MD system gray matter volume demonstrates a significant non-linear decreasing trend across childhood and adolescence ($p < .001$, see Figure S6a), exactly the same as that of the MD network ($p < .001$) used in the main text (see Figure S6b).

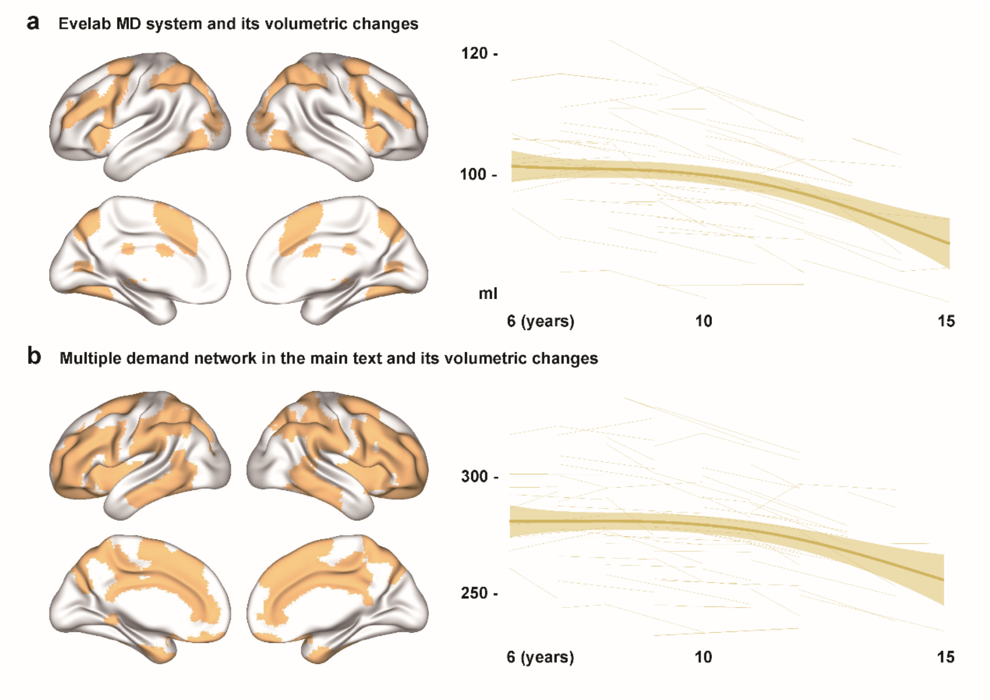


**Fig. S6 Comparisons between the two MD Networks**.

*Note*. Panel a demonstrates the Evelab MD system and its volumetric changes in the full sample. Panel b depicts the large-scale MD network discussed in the main text and its associated volumetric changes across the full sample.

TVEM analysis conducted for the gray matter volume in the Evelab MD system revealed a similar association pattern with overall language competence as observed in the MD network defined in the main text. We found significant positive associations between gray matter volume in the Evelab MD system and the Verbal Comprehension Index Vocabulary score in early school-age children (ages six to eight) and adolescents (ages 12 to 15). For the Similarities subtest, a significant positive association was also noted in school age children (ages six to eight), consistent with findings reported in the main text for the MD network. In addition, we found a significant positive association between the Comprehension subtest score and gray matter volume in the Evelab MD system in adolescents (ages 13 to 15), as shown in Figure S7a. These results support our original hypothesis about the two critical developmental periods where network-level gray matter volumes promote further advancement in overall language competence.

GAM estimates confirmed significant positive associations between gray matter volume in the Evelab MD system and both the Verbal Comprehension Index and Vocabulary score in early school-age children and adolescents, consistent with findings reported in the main text, as shown in Figure S7b and Table S3. Our results showed a marginally significant positive association between MD volume and the Similarities subtest score in children aged six to eight, aligning with the pattern reported in the main text. However, a potential link between the Comprehension subtest and the MD regions during adolescence was found (see Table S3), differing from the main text results with the large-scale MD network. This divergent finding suggests that the choice of analytical mask may affect the association between the Comprehension subtest and brain networks, indicating potential variability. Nonetheless, similar patterns as reported in the main text were observed using the Evelab MD system, reinforcing the significance of domain-general systems during critical developmental periods for further enhancement in overall language competence.

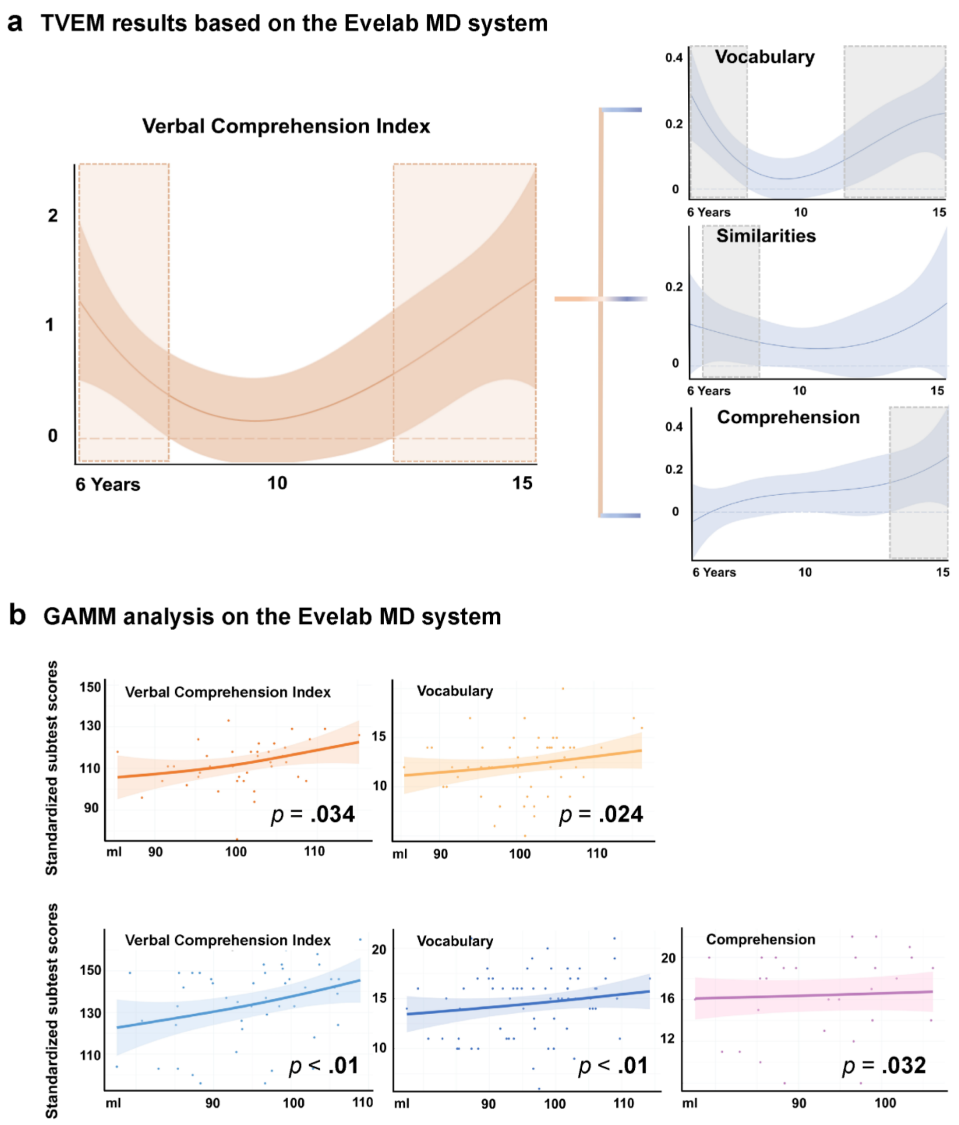


**Fig. S7 Age-related Associations between the Evelab MD System and Language Development.**

*Note.* Panel a depicts age-related associations between the Evelab MD system and the Verbal Comprehension Index (red) and its subtests (blue). Shaded areas indicate upper and lower 95% confidence intervals. Statistically significant associations ($p < .05$) between gray matter volume and language competence are indicated by the criterion dotted line falling out of the 95% confidence intervals. Dashed boxes highlight ages at which significant positive associations were observed between gray matter volume in the Evelab MD system and language development (as indicated by the fact that the 95% confidence band is above the estimated association of 0 on the $y$-axis). Panel b displays gray matter volume in the Evelab MD system associated with the Verbal Comprehension Index in specific age groups.

**Table S3**

GAM Estimates for Language Development Associated with the Evelab MD System.

| Group | Test | Best Model Fit | $R^2$ (adjusted) | Slope[1] | | | |
|---|---|---|---|---|---|---|---|
| | | | | edf | Red.df | F | $p$-value |
| Age 6-8 | VCI[2] | s(GMV[6])+sex | .128 | 1.654 | 2.063 | 3.703 | **.034** |
| | VOC[3] | s(GMV)+sex | .134 | 2.219 | 2.737 | 3.492 | **.024** |
| | SIM[4] | s(GMV)+sex | .105 | 2.231 | 2.762 | 2.371 | .079. |
| Age 12(13)-15 | VCI | s(GMV)+sex | .255 | 1.549 | 1.910 | 9.477 | <**.01** |
| | VOC | s(GMV)+sex | .084 | 1.000 | 1.001 | 7.689 | <**.01** |
| | COM[5] | s(GMV)+sex | .151 | 1.000 | 1.000 | 5.142 | **.032** |

[1]Smooth function (edf) as well as degrees of freedom (Red. df) and $F$-statistic and associated $p$-value for gray matter volume (**bold** highlights $p < .05$).

[2]VCI indicates the Verbal Comprehension Index.

[3]VOC denotes the Vocabulary subtest score.

[4]SIM represents the Similarities subtest score.

[5]COM refers to the Comprehension subtest score.

[6]GMV indicates gray matter volume in the Evelab MD system.

RWA was also conducted to estimate the unique contributions of the language network, the Evelab MD system, and the DMN to the further enhancement of the Verbal Comprehension Index and its subtest scores resulting from volumetric variations (see Figure S8 and Table S4). Results indicated that, similar to the MD network defined in the main text, the Evelab MD system made the most substantial contribution to overall language development due to network-level volumetric changes, accounting for over 44.67% of the increase in the Verbal Comprehension Index in children aged six to eight and over 61.61% in adolescents. For the subtests, gray matter volume in the language network was again the most important contributing factor during childhood, explaining more than 42% of the enhancement in subtest scores (the Vocabulary subtest: 39.68%; the Similarities subtest: 44.99%) attributed to volumetric variations. Moreover, while the gray matter volume in the DMN appeared to be the primary predictor of enhancement in Vocabulary score among adolescents (explaining more than 50% of the variance), relative weights revealed that the contributions of the Evelab

MD system and the DM network were basically equal (the Evelab MD system: .032; the DM network: .032).

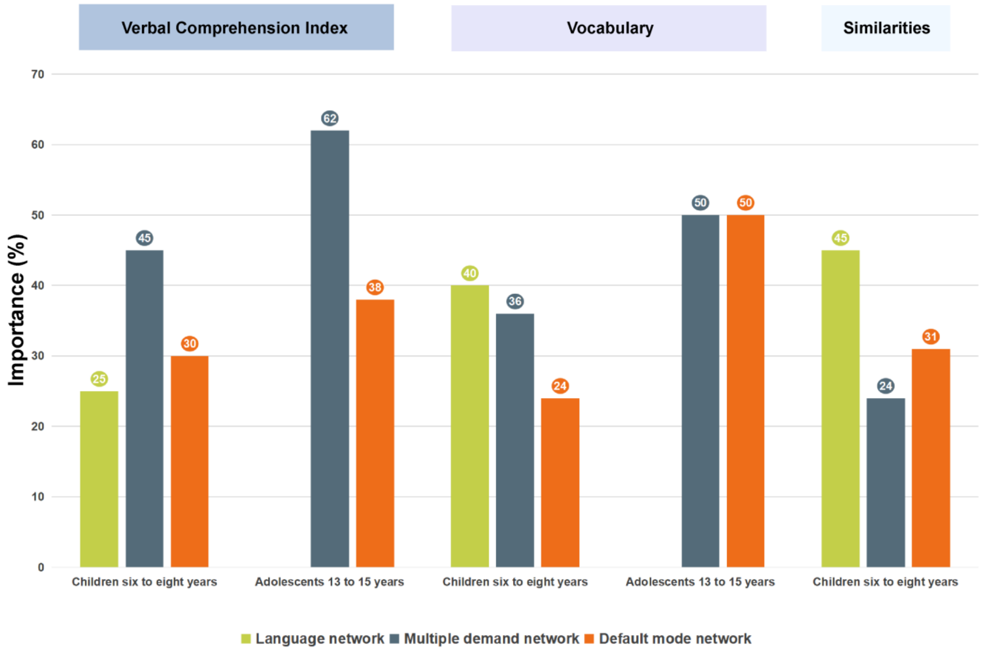


**Fig. S8 Network-level Volumetric Contributions to Language Development.**

**Table S4**

Relative Weight Analysis of Network GMV Predicting Overall Language Performance

| Network | Group | Relative Weights[1] | Importance (%)[2] | $\delta$ | Rank |
|---|---|---|---|---|---|
| Dependent Variable: the Verbal Comprehension Index | | | | | |
| LAN[3] | Age 6-8 | .031 | 25.17 | .122 | 3 |
| EveMD[4] | | .055 | **44.67**[6] | .280 | 1 |
| DM[5] | | .037 | 30.16 | .174 | 2 |
| $R^2 = .12$ | | | | | |
| EveMD | Age 13-15 | .178 | **61.61** | .517 | 1 |
| DM | | .111 | 38.39 | .149 | 2 |
| $R^2 = .29$ | | | | | |
| Dependent Variable: the Vocabulary Subtest Score | | | | | |
| LAN | Age 6-7 | .072 | **39.68** | .307 | 1 |
| EveMD | | .066 | 35.94 | .277 | 2 |
| DM | | .045 | 24.37 | .107 | 3 |
| $R^2 = .18$ | | | | | |
| EveMD | Age 13-15 | .032 | 49.73 | .177 | 2 |
| DM | | .032 | **50.27** | .180 | 1 |
| $R^2 = .06$ | | | | | |
| Dependent Variable: the Similarities Subtest Score | | | | | |
| LAN | Age 6-9 | .044 | **44.99** | .261 | 1 |
| EveMD | | .024 | 24.25 | .080 | 3 |
| DM | | .030 | 30.76 | .154 | 2 |
| $R^2 = .10$ | | | | | |

[1]Relative Weights is an estimated $R^2$ associated with each volumetric predictor.

[2]Importance (%): rescaled relative weights (relative weights divided by full model $R^2$).

[3]LAN: the language-selective network.

[4]EveMD: the Evelab MD system.

[5]DM: the default mode network.

[6]**Bold** indicates the strongest estimated predictor.

#### D Results Based on the Limbic System as the Control Network

The limbic system is frequently taken to underlie emotion and motivation (MacLean, 1952; Rolls, 2015). In order to investigate the specificity of our main findings for naturalistic language processing associated with the language-selective network and the two domain-general cognitive networks, we performed the same analyses for the control network we selected according to Yeo et al (2011), namely the limbic system (see Figure S9). First, the GAMM-based estimates revealed the same significant non-linear decreasing trend for the development of gray matter volume in the control limbic system ($p < .01$, see Figure S9).

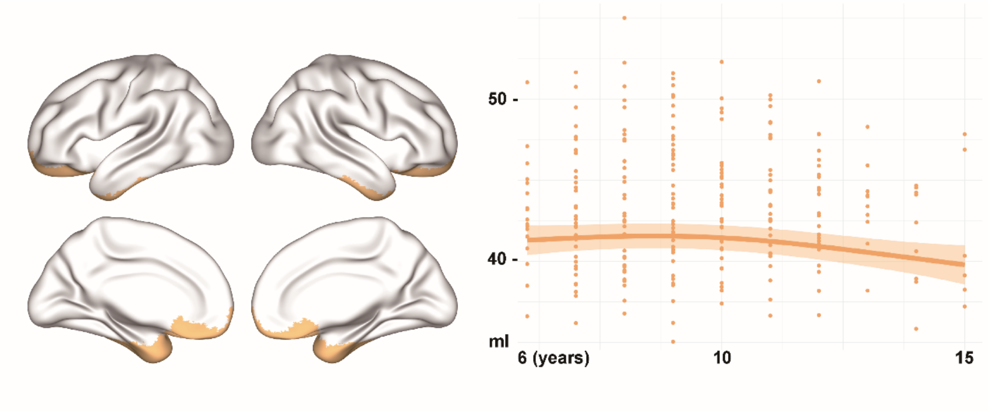


**Fig. S9 The Limbic System and its GMV Changes**.

Second, we performed the same TVEM analysis on the limbic system and found almost the same subgroups with relatively weak correlation between gray matter volume in the control network and overall language competence, as shown in Figure S10. However, GAM estimates further revealed that neither the competence of early school-age children between six and seven years old (the Verbal Comprehension Index: $p = .178$, the Similarities subtest: $p = .055$) nor that of adolescents aged 13 to 15 years ($p = .324$) significantly associated with the development of gray matter volume in the limbic system, see Table S5. The above results suggest a language-specific structural development across childhood and adolescence in both the language-selective network and the domain-general cognitive network, as opposed to the limbic system, which is not directly related to language processing.

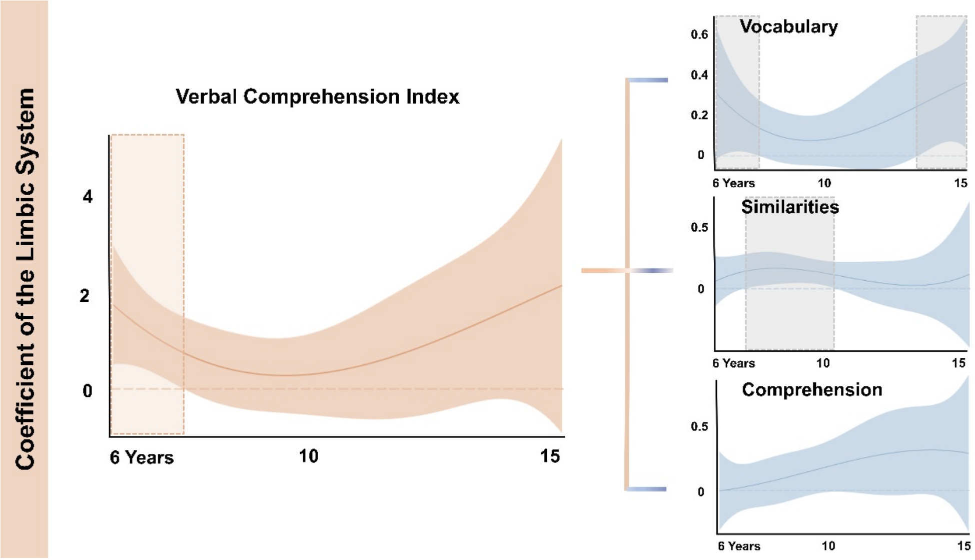


**Fig. S10 Age-related Associations of the Limbic System and Language Development.**

**Table S5**

GAM Estimates for Language Development Associated with the Evelab MD System.

| Group | Test | Best Model Fit | $R^2$ (adjusted) | Slope[1] | | | |
|---|---|---|---|---|---|---|---|
| | | | | edf | Red.df | F | $p$-value |
| Age 6-7 | VCI[2] | s(GMV[5])+sex | .055 | 1.000 | 1.000 | 1.924 | .178 |
| | VOC[3] | s(GMV)+sex | .192 | 1.000 | 1.000 | 7.480 | **.012** |
| Age 7-10 | SIM[4] | s(GMV)+sex | .073 | 1.001 | 1.002 | 3.752 | .055 |
| Age 13-15 | VOC | s(GMV) | .001 | 1.000 | 1.000 | 1.026 | .324 |

[1]Smooth function (edf) as well as degrees of freedom (Red. df) and $F$-statistic and associated $p$-value for gray matter volume (**bold** highlights $p < .05$).

[2]VCI indicates the Verbal Comprehension Index.

[3]VOC represents the Vocabulary subtest score.

[4]SIM denotes the Similarities subtest score.

[5]GMV indicates gray matter volume in the limbic system.

### E Results Based on the Purer Language Task of Audiovisual Integration

To compare the findings on overall language competence measured by the Wechsler Intelligence Scale in the main text with those obtained from lab-based research, we performed this exploratory analysis on an audiovisual integration task conducted in a laboratory setting. After visual inspection and data exclusion, a total of 256 scans from 200 participants (94 females) in the devCCNP-PEK Sample were included in this parallel analysis using the audiovisual integration task. Participants were required to judge whether the visually presented characters matched the auditory presented sounds. The stimulus set consisted of 40 Chinese characters, half of which had congruent audio-visual mappings and the other half had incongruent audio-visual mappings (see Figure S11). The procedure for the audiovisual integration task began with an 800 ms fixation cross, followed by the stimuli presented continuously until the participant responded, and ended with a blank screen before the onset of the next trial. The auditory and visual stimuli were initiated simultaneously. Participants were instructed to press the "J" key for congruent trials and the "F" key for incongruent trials. To avoid the speed-accuracy trade-off problem, we used the average ratio of reaction time to accuracy as the measure of performance for this task.

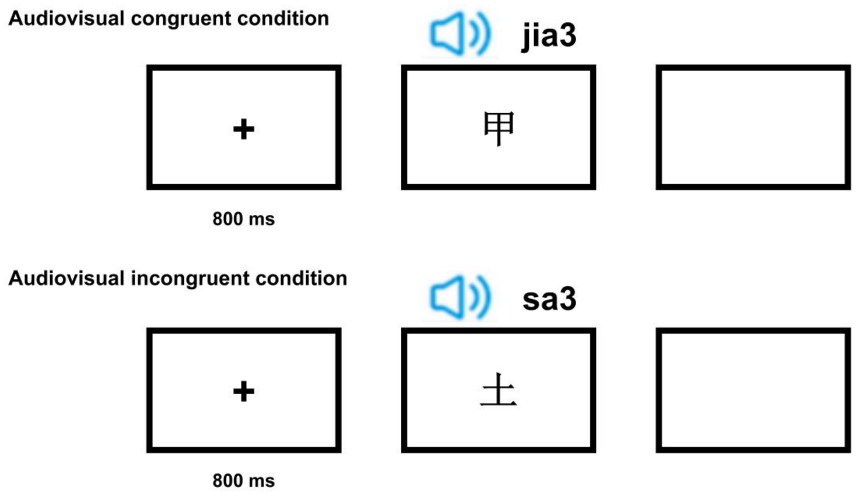


**Fig. S11 Experimental Procedure of the Purer Language Task Measuring Audiovisual Integration**.

To facilitate direct comparison with the results of verbal intelligence from the devCCNP-PEK Sample, we again conducted analyses specifically within the two predefined age windows: the early school-age period (6-8 years) and adolescence (13-15 years). In early school-age children, GAM estimates revealed marginally significant correlations between the average ratio and gray matter volumes in the language network ($p$ = .066) and the MD network ($p$ = .066), but not in the DM network ($p$ = .563). RWA indicated that the language network was the primary predictor during this period (relative weights: .02, importance: 65%), surpassing the MD network (relative weights: .01, importance: 35%).

In adolescence, GAM estimates revealed marginally significant correlations between the average ratio and the gray matter volume in the language network before correction ($p$ = .053), with no significant associations identified with the two domain-general cognitive networks (the MD network: $p$ = .072; the DM network: $p$ = .201).

For further details, please refer to Table S6. The results provide further evidence for a dissociation between core language functions and language abilities associated with general intelligence.

**Table S6**

GAM Estimates for Audiovisual Integration and its Volumetric Correlates.

| Group[1] | Network | Best Model Fit | $R^2$ (adjusted) | Slope[2] | | | |
|---|---|---|---|---|---|---|---|
| | | | | edf | Red.df | F | *p*-value |
| Age 6-8 | LAN[3] | s(GMV[5])+sex | .148 | 3.432 | 3.805 | 4.17 | .066(**.02** before correction) |
| | MD[4] | s(GMV)+sex | .124 | 3.249 | 3.702 | 3.49 | .066(**.04** before correction) |
| Age 13-15 | LAN | s(GMV)+sex | .160 | 1.498 | 1.826 | 3.02 | .108(**.05** before correction) |

[1]GAM estimates for gray matter volumes in the three networks associated with the audiovisual integration task in the devCCNP-PEK Sample.

[2]Smooth function (edf) as well as degrees of freedom (Red. df) and $F$-statistic and associated $p$-value for gray matter volumes (**bold** highlights $p < .05$).

[3]LAN: the language network.

[4]MD: the multiple demand network.

[5]GMV indicates network-level gray matter volumes.

### F Results for Hemispheric Analysis

**Table S7**

GAMM Estimates for Hemispheric GMV in the Three Networks Separately.

| Hemisphere | Best Model Fit | $R^2$ (adjusted) | Age Spline[1] | | | |
|---|---|---|---|---|---|---|
| | | | edf | Red.df | F | *p*-value |
| L[2]LAN[4] | s(age)+sex | .247 | 2.85 | 2.85 | 12.35 | <**.001** |
| R[3]LAN | s(age)+sex | .255 | 2.66 | 2.66 | 13.26 | <**.001** |
| LMD[5] | s(age)+sex | .266 | 2.82 | 2.82 | 27.12 | <**.001** |
| RMD | s(age)+sex | .259 | 2.74 | 2.74 | 29.55 | <**.001** |
| LDM[6] | s(age)+sex | .258 | 2.73 | 2.73 | 15.04 | <**.001** |
| RDM | s(age)+sex | .267 | 2.87 | 2.87 | 20.43 | <**.001** |

[1]Smooth function (edf) as well as degrees of freedom (Red. df) and $F$-statistic and associated $p$-value for gray matter volume (**bold** highlights $p < .05$).

[2]L indicates the left hemisphere.

[3]R indicates the right hemisphere.

[4]LAN: the gray matter volume in the language-selective network.

[5]MD: the gray matter volume in the multiple demand network.

[6]DM: the gray matter volume in the default mode network.

**Table S8**

GAM Estimates for Hemispheric GMV Associated with the Verbal Comprehension Index across Age Groups.

| Group | Hemisphere | Best Model Fit | $R^2$ (adjusted) | Slope[1] | | | |
|---|---|---|---|---|---|---|---|
| | | | | edf | Red.df | F | $p$-value |
| Age 6-8 | L[2]LAN[4] | s(GMV[7])+sex | .07 | 1.000 | 1.000 | 4.678 | < **.05** |
| | R[3]LAN | s(GMV)+sex | .01 | 1.000 | 1.000 | 2.345 | .135 |
| | LMD[5] | s(GMV)+sex | .17 | 1.515 | 1.861 | 5.645 | < **.05** |
| | RMD | s(GMV)+sex | .14 | 1.003 | 1.006 | 7.795 | < **.01** |
| | LDM[6] | s(GMV)+sex | .12 | 1.001 | 1.001 | 6.921 | < **.05** |
| | RDM | s(GMV)+sex | .05 | 1.515 | 1.285 | 2.595 | .0821 |
| Age 13-15 | LMD | s(GMV)+sex | .33 | 1.000 | 1.000 | 10.30 | < **.01** |
| | RMD | s(GMV)+sex | .26 | 1.000 | 1.000 | 8.03 | < **.05** |
| | LDM | s(GMV)+sex | .26 | 1.000 | 1.000 | 7.83 | < **.05** |
| | RDM | s(GMV)+sex | .27 | 1.000 | 1.000 | 8.32 | < **.05** |

[1]Smooth function (edf) as well as degrees of freedom (Red. df) and $F$-statistic and associated $p$-value for gray matter volume (**bold** highlights $p < .05$).

[2]L indicates the left hemisphere.

[3]R indicates the right hemisphere.

[4]LAN: the gray matter volume in the language-selective network.

[5]MD: the gray matter volume in the multiple demand network.

[6]DM: the gray matter volume in the default mode network.

[7]GMV indicates network-level gray matter volumes.

**Table S9**

GAM Estimates for Hemispheric GMV Associated with the Vocabulary Subtest across Age Groups.

| Group | Hemisphere | Best Model Fit | $R^2$ (adjusted) | Slope[1] | | | |
|---|---|---|---|---|---|---|---|
| | | | | edf | Red.df | F | $p$-value |
| Age 6-7 | L[2]LAN[4] | s(GMV[7])+sex | .193 | 1.000 | 1.001 | 6.971 | $<$ **.05** |
| | R[3]LAN | s(GMV)+sex | .187 | 1.030 | 1.059 | 6.226 | $<$ **.05** |
| | LMD[5] | s(GMV)+sex | .243 | 1.590 | 1.963 | 4.933 | $<$ **.05** |
| | RMD | s(GMV)+sex | .189 | 1.057 | 1.112 | 6.369 | $<$ **.05** |
| | LDM[6] | s(GMV)+sex | .193 | 1.000 | 1.000 | 6.974 | $<$ **.05** |
| | RDM | s(GMV)+sex | .175 | 1.000 | 1.000 | 6.326 | $<$ **.05** |
| Age 13-15 | LMD | s(GMV)+sex | .216 | 1.000 | 1.000 | 5.412 | $<$ **.05** |
| | RMD | s(GMV)+sex | .207 | 1.000 | 1.000 | 5.139 | $<$ **.05** |
| | LDM | s(GMV)+sex | .197 | 1.000 | 1.000 | 4.811 | $<$ **.05** |
| | RDM | s(GMV)+sex | .246 | 1.000 | 1.000 | 6.42 | $<$ **.05** |

[1]Smooth function (edf) as well as degrees of freedom (Red. df) and $F$-statistic and associated $p$-value for gray matter volume (**bold** highlights $p < .05$).

[2]L indicates the left hemisphere.

[3]R indicates the right hemisphere.

[4]LAN: the gray matter volume in the language-selective network.

[5]MD: the gray matter volume in the multiple demand network.

[6]DM: the gray matter volume in the default mode network.

[7]GMV indicates network-level gray matter volumes.

**Table S10**

GAMM Estimates for Hemispheric GMV Associated with the Similarities Subtest across Age Group.

| Group | Hemisphere | Best Model Fit | $R^2$ (adjusted) | Slope[1] | | | |
|---|---|---|---|---|---|---|---|
| | | | | edf | Red.df | F | $p$-value |
| Age 6-10 | L[2]LAN[4] | s(GMV[7]) | .0690 | 1.000 | 1.000 | 6.681 | < **.05** |
| | R[3]LAN | s(GMV) | .1040 | 1.000 | 1.000 | 9.408 | < **.05** |
| | LMD[5] | s(GMV) | .0892 | 1.000 | 1.000 | 2.942 | < **.05** |
| | RMD | s(GMV) | .0690 | 1.000 | 1.000 | 6.033 | < **.05** |
| | LDM[6] | s(GMV) | .0681 | 1.000 | 1.000 | 6.465 | < **.05** |
| | RDM | s(GMV) | .0671 | 1.000 | 1.000 | 5.932 | < **.05** |

[1]Smooth function (edf) as well as degrees of freedom (Red. df) and $F$-statistic and associated $p$-value for gray matter volume (**bold** highlights $p < .05$).

[2]L indicates the left hemisphere.

[3]R indicates the right hemisphere.

[4]LAN: the gray matter volume in the language-selective network.

[5]MD: the gray matter volume in the multiple demand network.

[6]DM: the gray matter volume in the default mode network.

[7]GMV indicates network-level gray matter volumes.

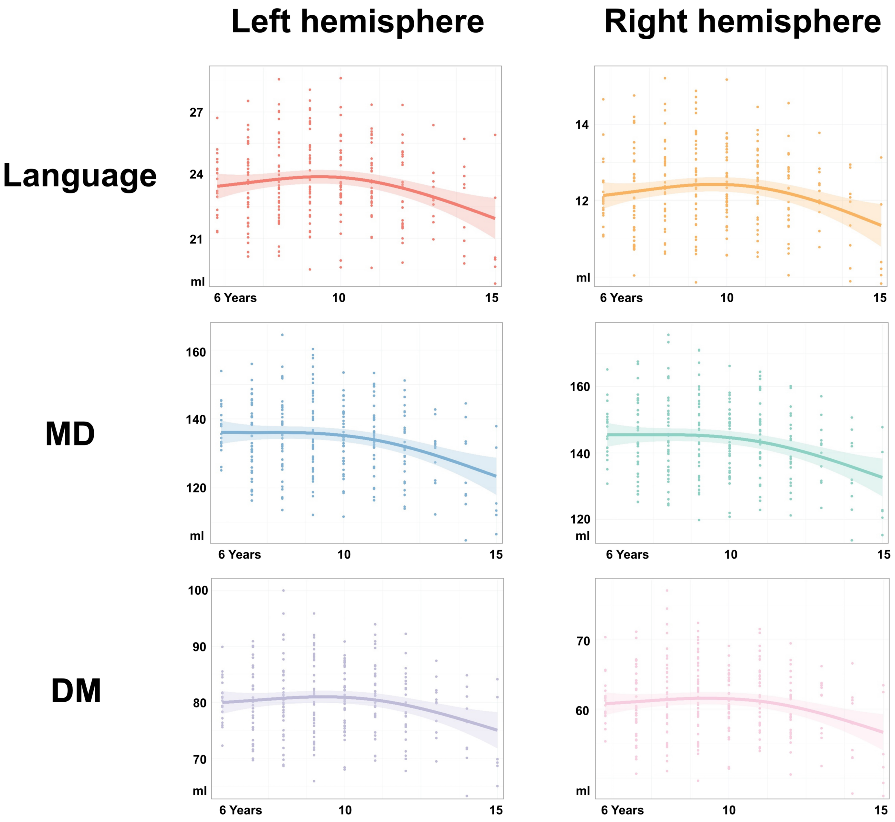


**Fig. S12** **Hemispheric Developmental Trajectories for GMV in the three Networks.**

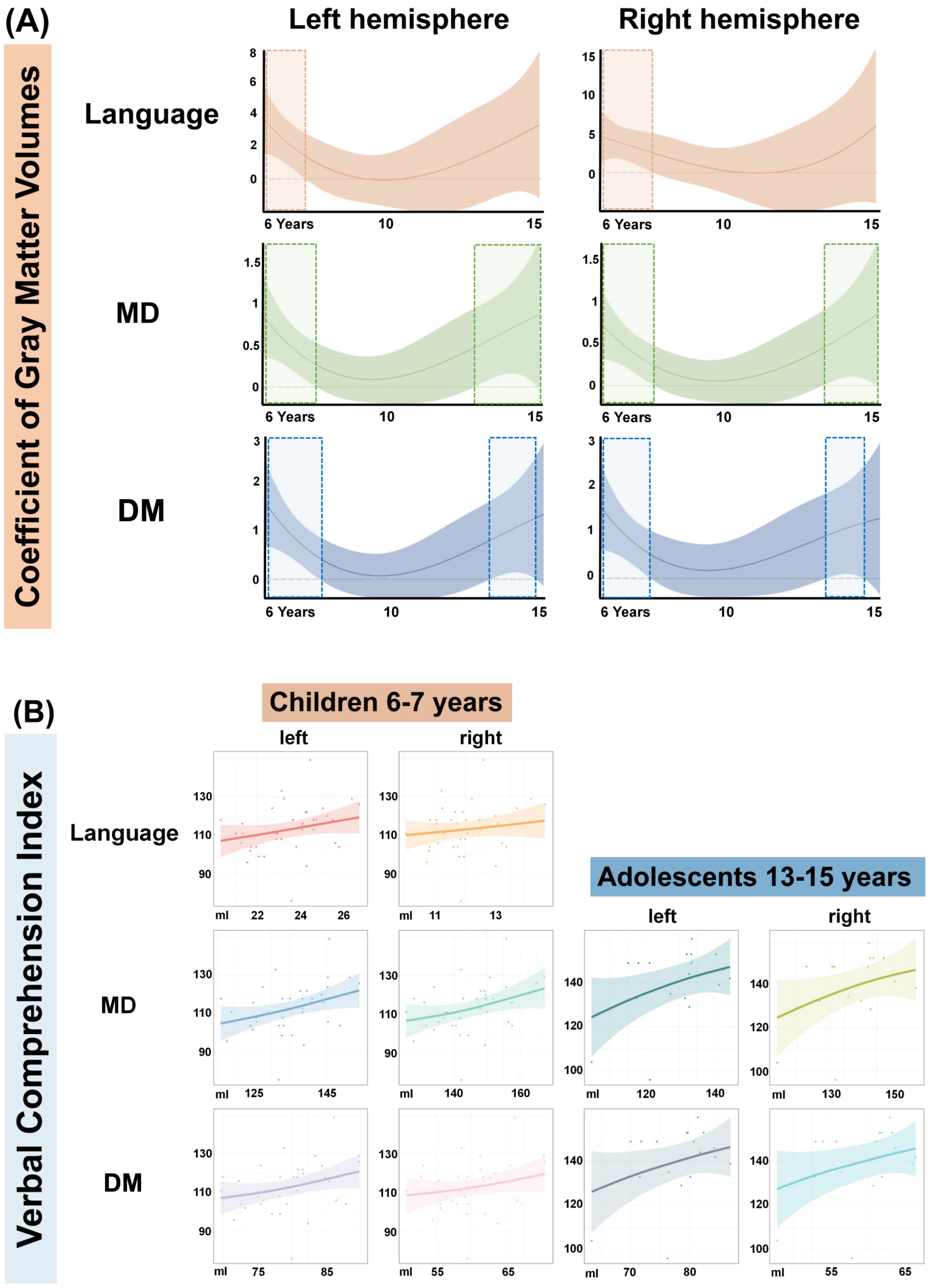


**Fig. S13 Association Curves between Hemispheric GMV and the Verbal Comprehension Index.**



*Note.* A indicates the full sample while B indicates subgroups of 6-8 and 13-15 years.

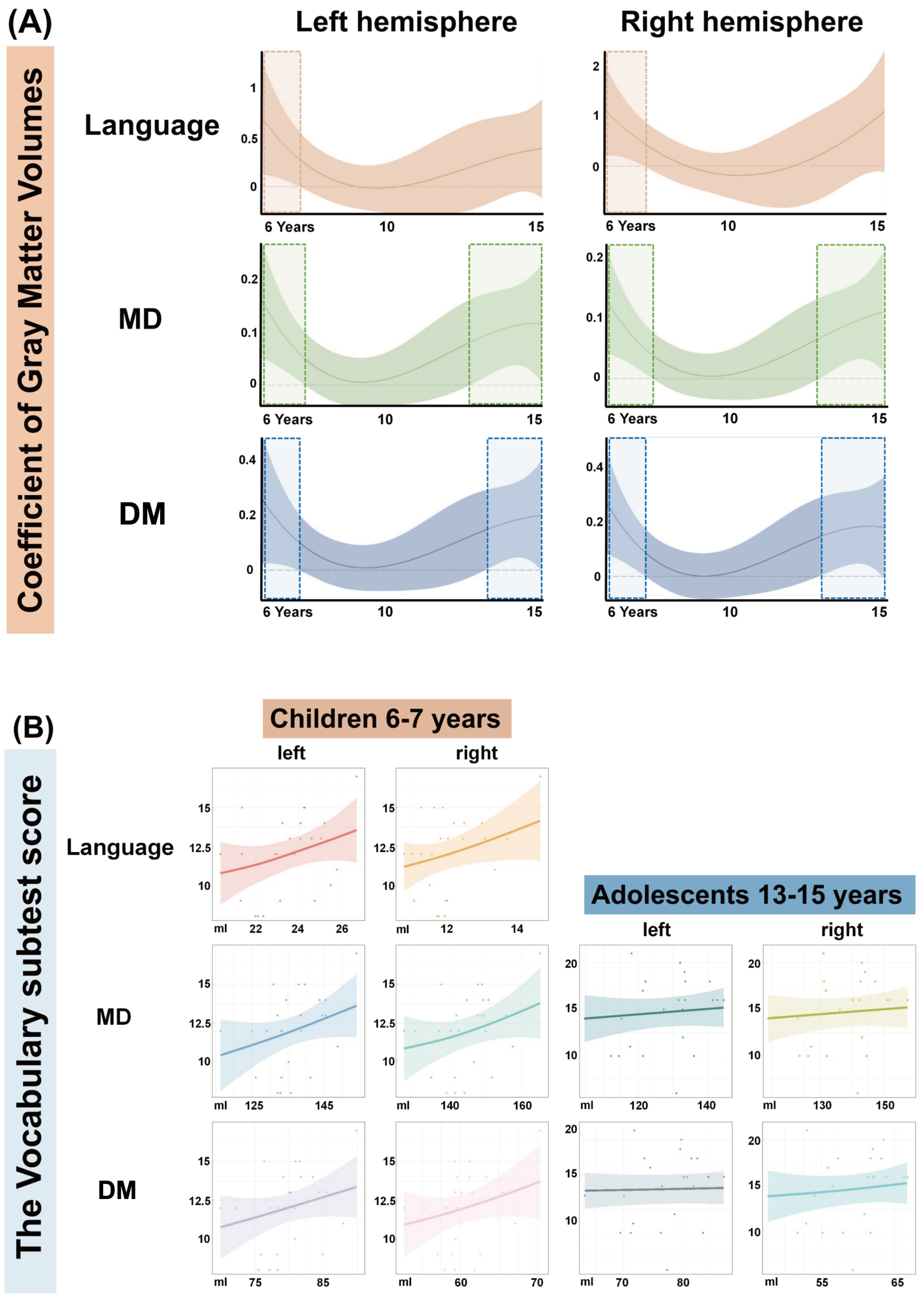


**Fig. S14 Association Curves between Hemispheric GMV and the Vocabulary Subtest.**

*Note.* A indicates the full sample while B indicates subgroups of 6-7 and 13-15 years.

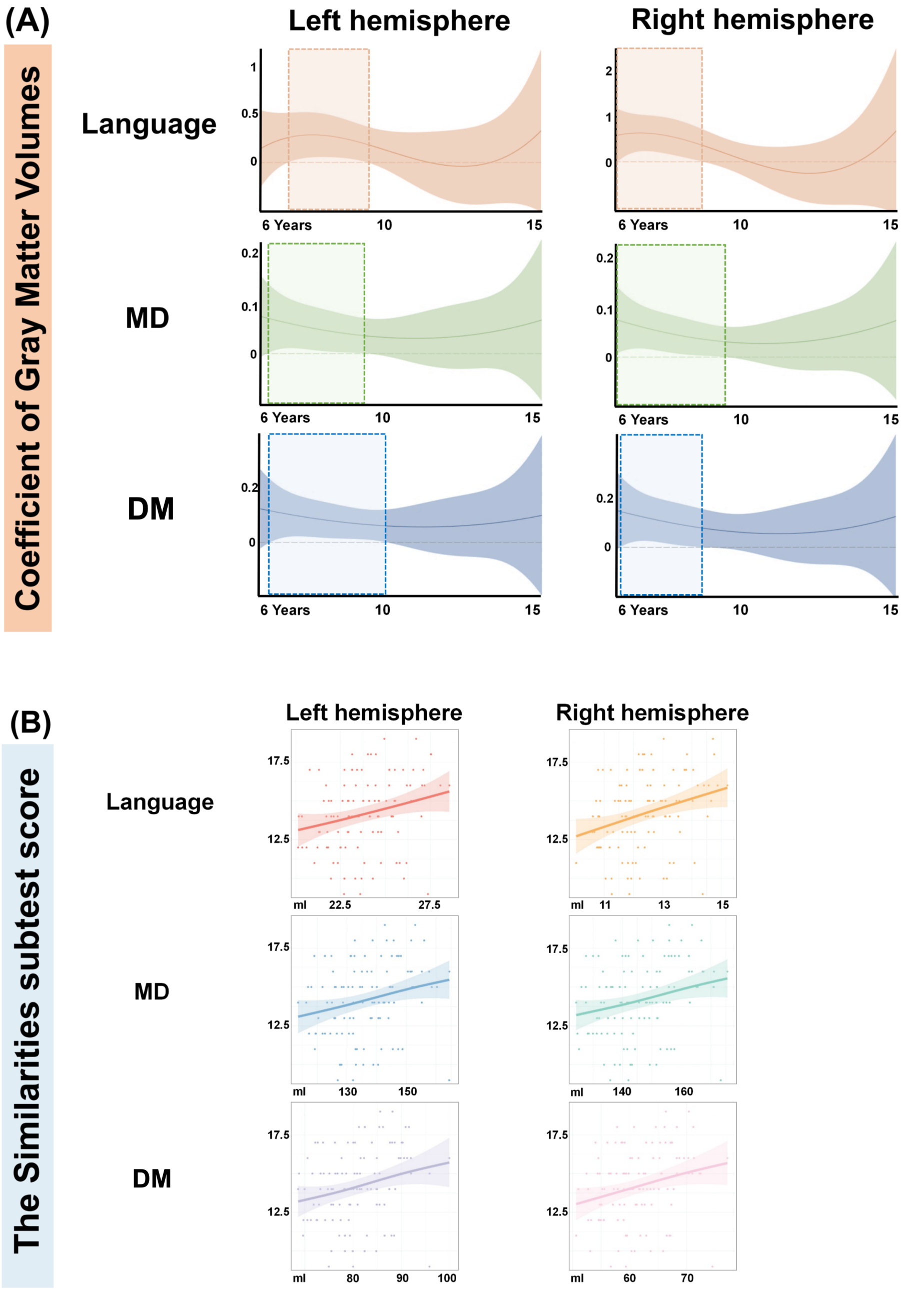


**Fig. S15 Association Curves between Hemispheric GMV and the Similarities Subtest.**

*Note.* A indicates the full sample while B indicates the subgroup of 6-10 years.

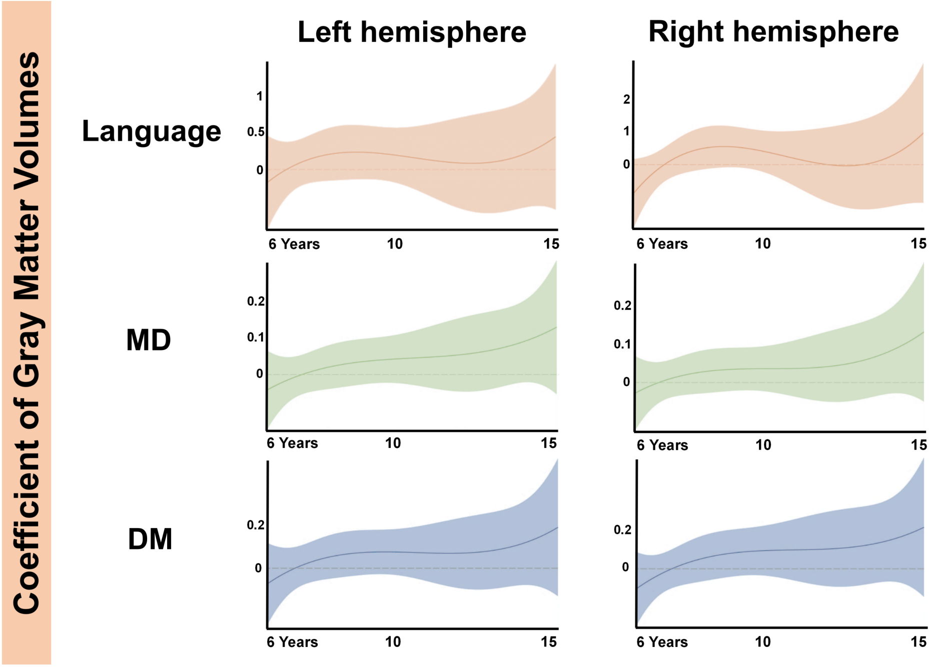


**Fig. S16** **Association Curves between Hemispheric GMV and the Comprehension Subtest.**
